# Purinergic signaling controls spontaneous activity in the auditory system throughout early development

**DOI:** 10.1101/2020.08.11.246306

**Authors:** Travis A. Babola, Sally Li, Zhirong Wang, Calvin Kersbergen, Ana Belén Elgoyhen, Thomas Coate, Dwight Bergles

## Abstract

Spontaneous bursts of electrical activity in the developing auditory system arise within the cochlea prior to hearing onset and propagate through future sound processing circuits of the brain to promote maturation of auditory neurons. Studies in isolated cochleae revealed that this intrinsically generated activity is initiated by ATP release from inner supporting cells (ISCs), resulting in activation of purinergic autoreceptors, K^+^ efflux and subsequent depolarization of inner hair cells (IHCs). However, little is known about when this activity emerges or whether different mechanisms underlie distinct stages of development. Here we show that spontaneous electrical activity in mouse cochlea emerges within ISCs during the late embryonic period, preceding the onset of spontaneous correlated activity in IHCs and spiral ganglion neurons (SGNs), which begins at birth and follows a base to apex developmental gradient. At all developmental stages, pharmacological inhibition of P2Y1 metabotropic purinergic receptors dramatically reduced spontaneous activity in these three cell types. Moreover, *in vivo* imaging within the inferior colliculus of awake mice revealed that auditory neurons within future isofrequency zones exhibit coordinated neural activity at birth. The frequency of these discrete bursts increased progressively during the postnatal prehearing period, yet remained dependent on P2RY1. Analysis of mice with disrupted cholinergic signaling in the cochlea, indicate that this input modulates, rather than initiates, spontaneous activity before hearing onset. Thus, the auditory system uses a consistent mechanism involving ATP release from ISCs and activation of purinergic autoreceptors to elicit coordinated excitation of neurons that will process similar frequencies of sound.

**SIGNIFICANCE STATEMENT:** In developing sensory systems, groups of neurons that will process information from similar sensory space exhibit highly correlated electrical activity that is critical for proper maturation and circuit refinement. Defining the period when this activity is present, the mechanisms responsible and the features of this activity are crucial for understanding how spontaneous activity influences circuit development. We show that, from birth to hearing onset, the auditory system relies on a consistent mechanism to elicit correlate firing of neurons that will process similar frequencies of sound. Targeted disruption of this activity will increase our understanding of how these early circuits mature and may provide insight into processes responsible for developmental disorders of the auditory system.

## INTRODUCTION

In the developing central nervous system, spontaneous bursts of electrical activity promote maturation of newly formed neural circuits by promoting cell specification, survival, and refinement (Blankenship and Feller, 2010). These periodic bouts of electrical activity are prominent in developing sensory systems, where they arise through sensory-independent mechanisms. In the visual system, intrinsically-generated bursts of electrical activity, termed retinal waves, sweep across the retina (Feller et al., 1996) and, when disrupted, lead to refinement deficits in higher visual centers (Rossi et al., 2001; Xu et al., 2011; Zhang et al., 2012). The mechanisms responsible for generating this activity are dynamic and progress through distinct stages before eye opening, with early waves mediated by gap-junction coupling and later by acetylcholine and glutamate release from starburst amacrine and bipolar cells, respectively (Firth et al., 2005; Blankenship and Feller, 2010). Across these stages, neural activity changes dramatically, progressing from individual propagating waves to multiple wavelets with complex patterns that can be modulated by external light that penetrates the eyelid (Tiriac et al., 2018; Gribizis et al., 2019). These results demonstrate that the visual system uses an intricate process to shift activity patterns according to developmental stage, which may be optimized to achieve distinct aspects of circuit maturation. Although the visual system provides a template to understand developmental changes in sensory pathways, it is not known if similar processes are used to promote refinement in other sensory systems, limiting our understanding of how these circuits use intrinsically generated activity to induce maturation and refinement.

In the developing auditory system, peripheral and central neurons exhibit periodic bursts of action potentials that originate within the cochlea (Lippe, 1994; Tritsch et al., 2007, 2010). Prior to hearing onset, a group of glial-like inner supporting cells (ISCs) located adjacent to inner hair cells (IHCs) spontaneously release ATP, activating a metabotropic purinergic cascade that ultimately results in release of K^+^, IHC depolarization and subsequent burst firing of spiral ganglion (SGNs) and central auditory neurons (Sonntag et al., 2009; Babola et al., 2018). Recent mechanistic studies revealed that activation of purinergic P2Y1 receptors and downstream gating of Ca^2+^-activated chloride channels (TMEM16A) are required, and that correlated activity in central auditory circuits is sensitive to P2RY1 inhibition *in vivo* after the first postnatal week (Wang et al., 2015; Babola et al., 2020). Efferent inhibition of IHCs through activation of α9 subunit-containing nicotinic acetylcholine receptors has also been implicated in both initiating (Johnson et al., 2012) and modulating (Clause et al., 2014) spontaneous activity during this period. Although SGNs can fire action potentials as early as E14.5 (Marrs and Spirou, 2012),, it is not known when burst firing begins within the cochlea or what specific mechanisms initiate this activity at each developmental stage. Understanding these processes may help define the discrete steps required for maturation of precise auditory circuits and enhance our understanding of developmental auditory disorders.

Here, we examined the mechanisms responsible for initiating spontaneous activity in embryonic and postnatal mouse cochleae prior to hearing onset. Our results indicate that ISC electrical activity requires release of ATP and activation of P2Y1 autoreceptors at all developmental stages. Consistent with the critical role of ISC activation in triggering periodic activation of IHCs and SGNs, acute pharmacological inhibition of P2RY1 disrupted correlated activation of IHCs and SGNs during this period. *In vivo* imaging of auditory midbrain neurons in neonatal awake mice revealed that neurons within future isofrequency zones exhibit correlated activity at birth, providing a two-week period of highly stereotyped activity with which to influence circuit maturation. The frequency of these events increased progressively over the first two postnatal weeks, but remained dependent on P2RY1. Together, these studies suggest that, in contrast to the developing visual system, the auditory system uses a persistent mechanism involving ISC ATP release and activation of purinergic autoreceptors to elicit periodic bursts of activity in discrete groups of sensory neurons that will process similar frequencies of sound.

## MATERIALS AND METHODS

Both male and female between embryonic day 14 (E14) and postnatal day 16 (P16) were used for all experiments and randomly allocated to experimental groups. All animals were healthy and were only used for experiments detailed in this study. Transgenic breeders were crossed to female FVB/NJ (Friend Virus B NIH Jackson; demonstrated low hearing thresholds at 28 weeks) mice to improve litter sizes and pup survival (Zheng et al., 1999). For these studies, all mouse lines were maintained on mixed backgrounds, except for *Snap25-T2A-GCaMP6s* mice used in Figure 7 and 8, which were maintained on a C57BL/6 background. Mice were housed on a 12 hour light/dark cycle and were provided food ad libitum. This study was performed in accordance with recommendations provided in the Guide for the Care and Use of Laboratory Animals of the National Institutes of Health. All experiments and procedures were approved by the Johns Hopkins Institutional Care and Use Committee (protocol #: M018M330) and the Georgetown University Institutional Animal Care and Use Committee (protocol #1147). Surgery was performed under isoflurane anesthesia and extensive effort was made to minimize animal suffering.

**Figure 1.**
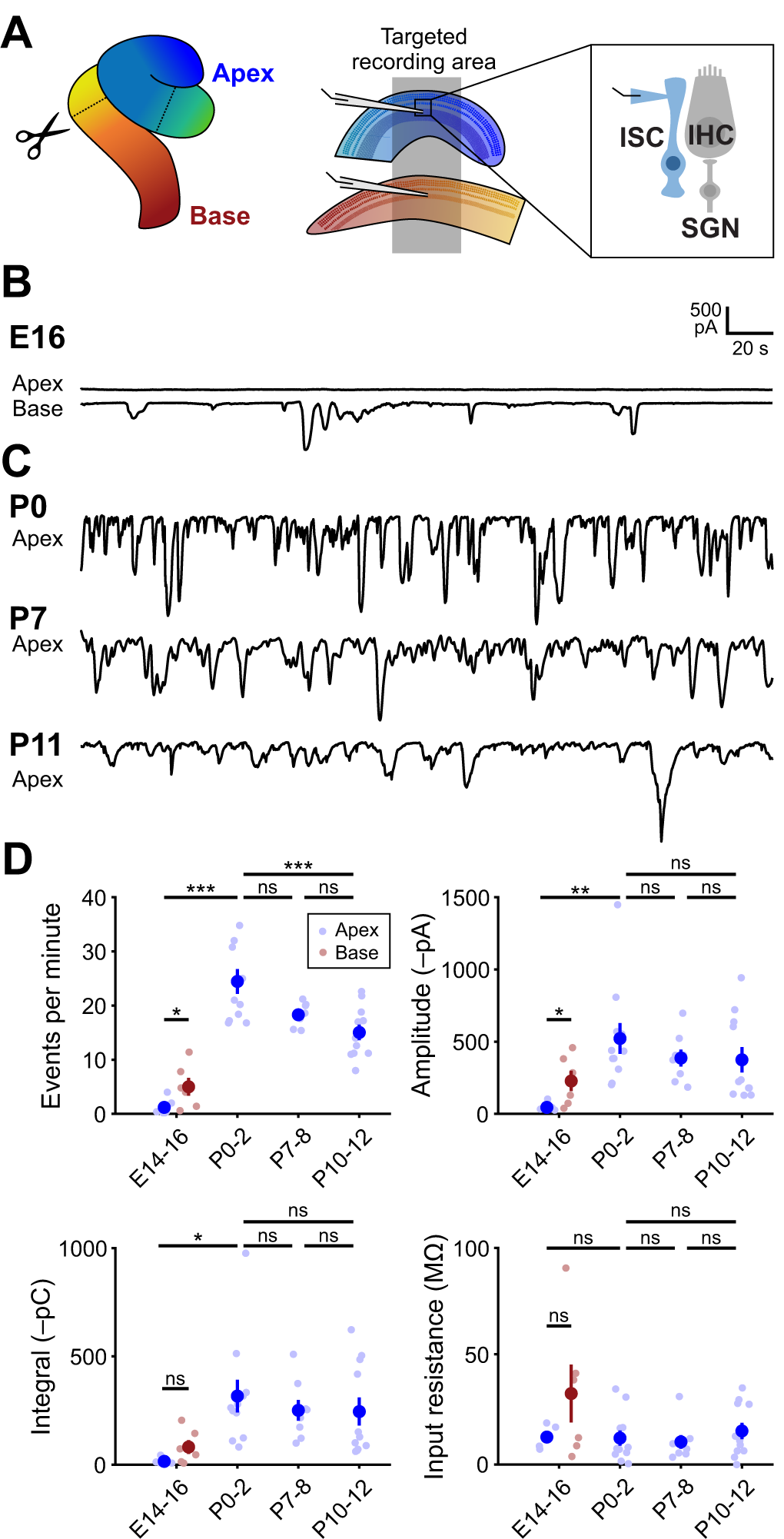
Prenatal onset of spontaneous activity in cochlear inner supporting cells. (A) Diagram of cochlea with approximate locations of cuts used for targeted apical and basal recordings of ISCs. (B) Schematic of whole-cell recording configuration from ISCs. Recordings were made at near physiological temperatures (32-34°C). (C) Exemplar voltage-clamp recordings of E16 apical and basal ISCs from the same cochlea. (D) Spontaneous inward currents in apical ISCs at different postnatal ages. (E) Quantification of ISC spontaneous current frequency, amplitude, integral (charge transfer), and input resistance. Data shown as mean ± SEM. n = 12 E14-16 recordings, 6 cochleae (6 apex and 6 paired base) from 6 mice, n = 11 P0-2 recordings 11 cochleae from 6 mice, n = 8 P7-8 recordings, 8 cochleae from 8 mice, and n = 11 P10-12 recordings, 11 cochleae from 8 mice. Comparisons between E16 apex and base, (Mann-Whitney U with Benjamini-Hochberg adjustment; *p < 0.05, ns: not significant). For comparisons between ages, E14-16 base and apex were combined into one group, (one-way ANOVA with Tukey post-hoc; ***p < 5e-4, **p < 0.005, *p < 0.05, ns: not significant).

**Figure 2.**
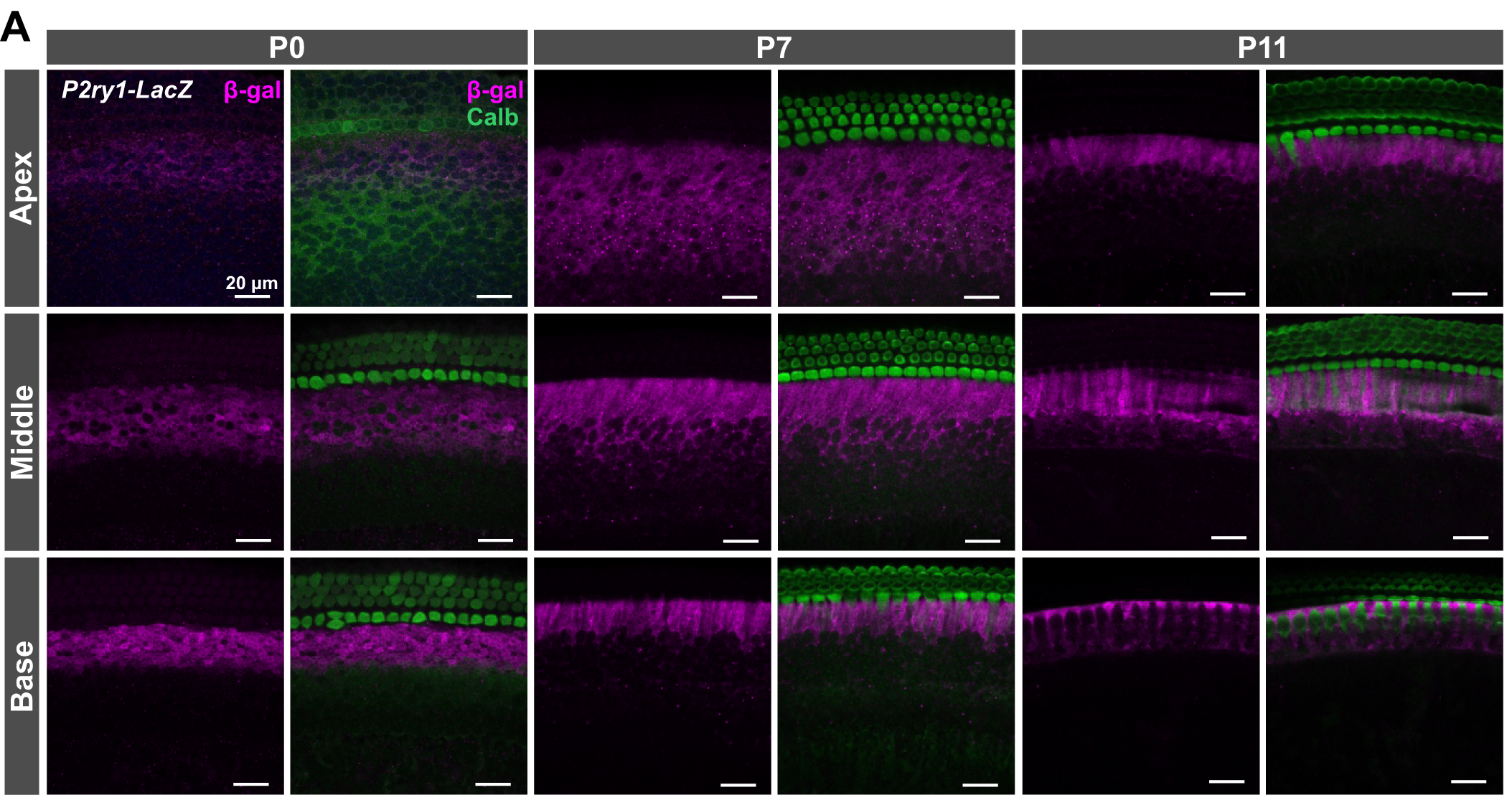
P2ry1 promoter activity in inner supporting cells throughout the prehearing period. (A) Immunostaining of cochlear sections from P0-P11 *P2ry1-LacZ* mice for B-galactosidase (rabbit anti-B-gal, Sanes lab) and calbindin (goat anti-calbindin, Santa Cruz) for labeling of hair cells.

**Figure 3.**
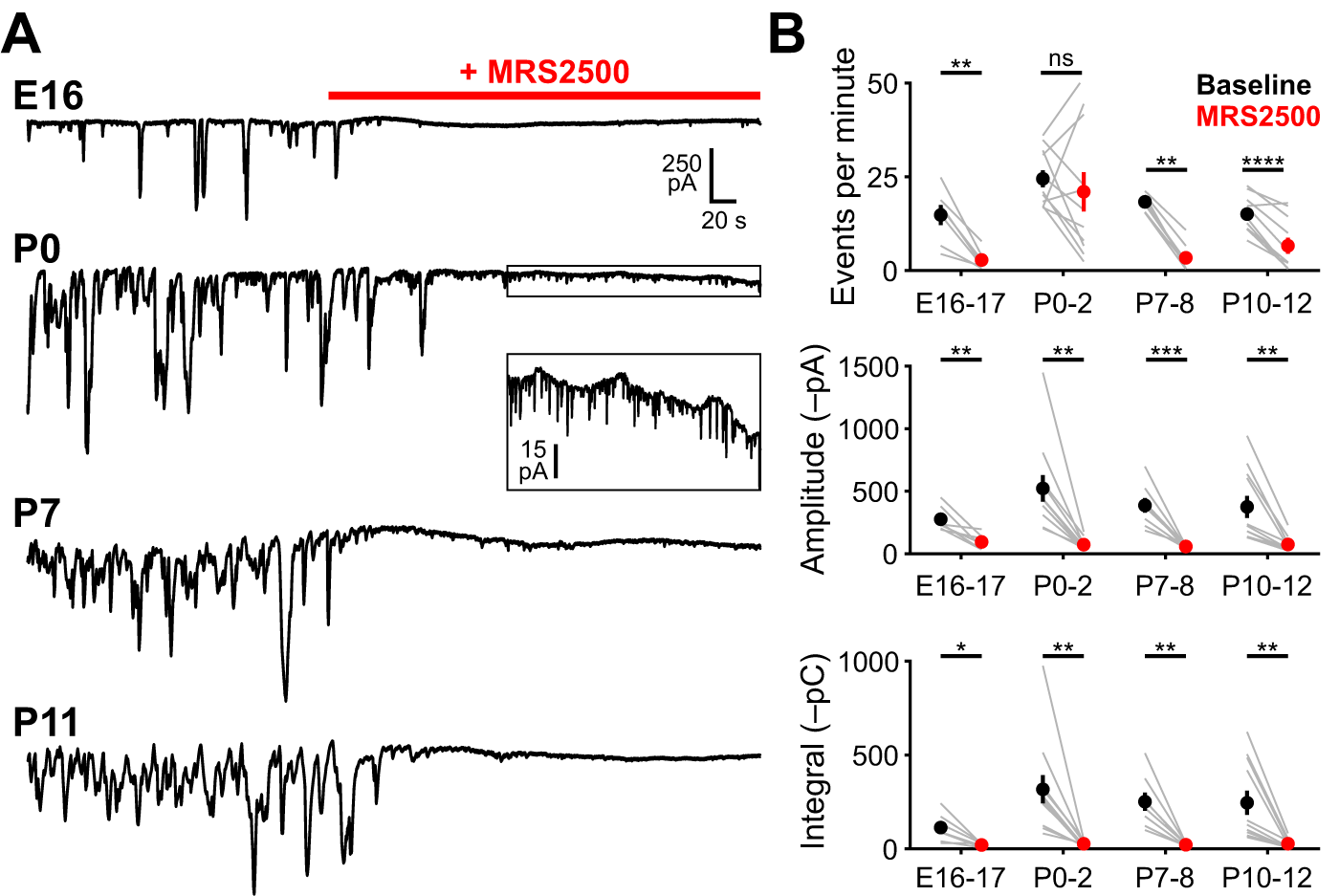
P2RY1 mediates supporting cell spontaneous currents in the cochlea throughout the prehearing period. (A) Spontaneous inward currents recorded from ISCs before and during application of MRS2500 (1 μM) at different developmental ages. Recordings were made at near physiological temperature (32-34°C). (B) Quantification of ISC spontaneous current frequency, amplitude, and integral (charge transfer) before and after application of MRS2500. n = 7 E16-17 ISCs, 7 cochleae from 7 mice, n = 11 P0-2 ISCs, 11 cochleae from 8 mice, n = 8 P7-8 ISCs, 8 cochleae from 8 mice, and n = 11 cochleae from 8 mice, (Students paired t-test with Bonferroni correction, ***p < 5e-4, **p < 0.005, *p < 0.05, ns: not significant).

**Figure 4.**
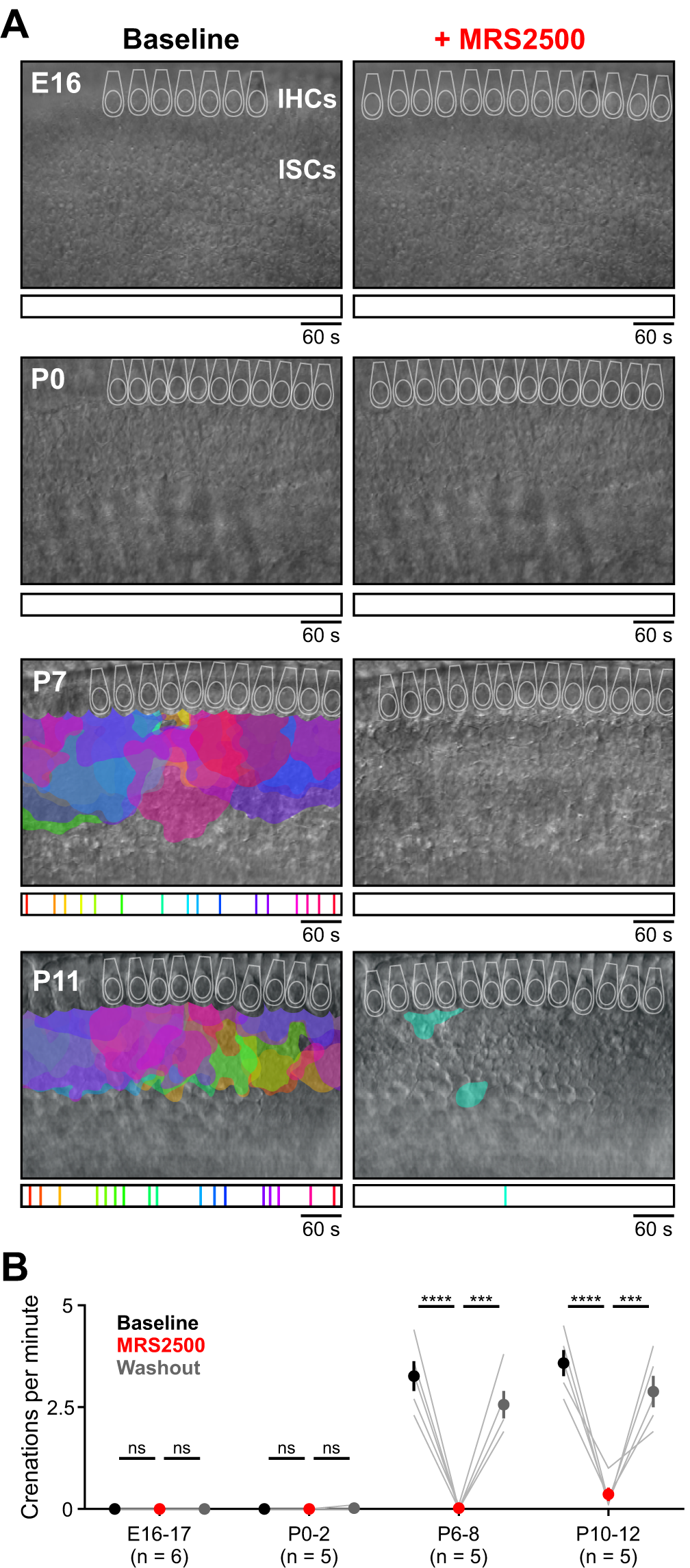
Delayed onset of P2ry1-dependent spontaneous crenations in ISCs. (A) Intrinsic optical imaging performed before and after application of the P2RY1 antagonist, MRS2500 (1 μM). Detected crenations are outlined in colors based on time of occurrence as indicated by timeline below image. Imaging was performed near physiological temperature (32-34°C). (B) Plot of crenation frequency and area before and after application of MRS2500. n = 6 E16-17 videos, 6 cochleae from 6 mice, n = 5 P0-2 videos, 5 cochleae from 5 mice, n = 5 P6-8 videos, 5 cochleae from 5 mine, and n = 5 P10-12 videos, 5 cochleae from 5 mice, (one-way ANOVA with Tukey post-hoc; ****p < 5e-5, ***p < 5e-4, ns: not significant).

**Figure 5.**
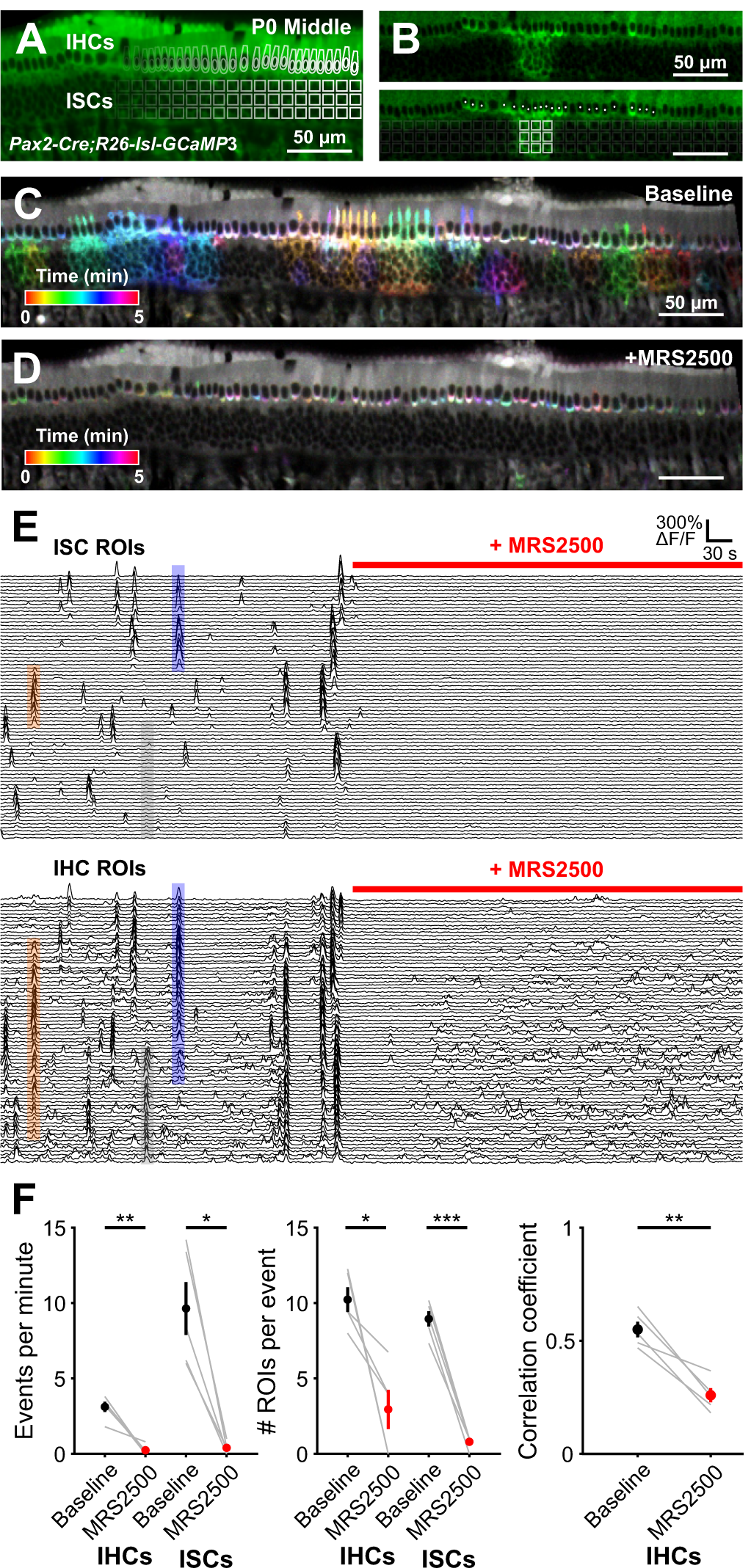
Correlated activation of IHCs and ISCs requires P2RY1 signaling. (A) Image of an excised cochlea from a P0 *Pax2-Cre;R26-lsl-GCaMP3* mouse. For analysis of time lapse imaging, a grid of square ROIs was placed over the ISCs and single ROIs were drawn for each IHC. Imaging was performed at near physiological temperature (32-34°C). (B) Exemplar Ca^2+^ transient in ISCs and simultaneous activation of multiple IHC. (bottom) Circles indicate active IHCs, white squares indicate active ISCs. (C) Ca^2+^ transients in control (baseline) conditions colored based on time of occurrence. (D) Ca^2+^ transients with P2RY1 inhibited (MRS2500, 1 μM) conditions colored based on time of occurrence. (E) Individual ROI traces for ISCs (top, 100 randomly selected) and IHCs (bottom, all shown). Colored boxes are examples of coordinated activity of ISCs and IHCs. Note that the number of IHCs activated can extend far beyond area of ISC activation (see Figure 6). Grey box indicates IHC activation on the edge of the frame with no ISC activation, likely caused by an out-of-frame ISC event. (F) Quantification of coordinated event frequency, number of ROIs per coordinated event, and the correlation coefficient before and after application of MRS2500. n = 5 cochleae from 3 mice, (paired t-test with Benjamini-Hochberg adjustment, ***p < 5e-4,**p < 0.005, *p < 0.05).

**Figure 6.**
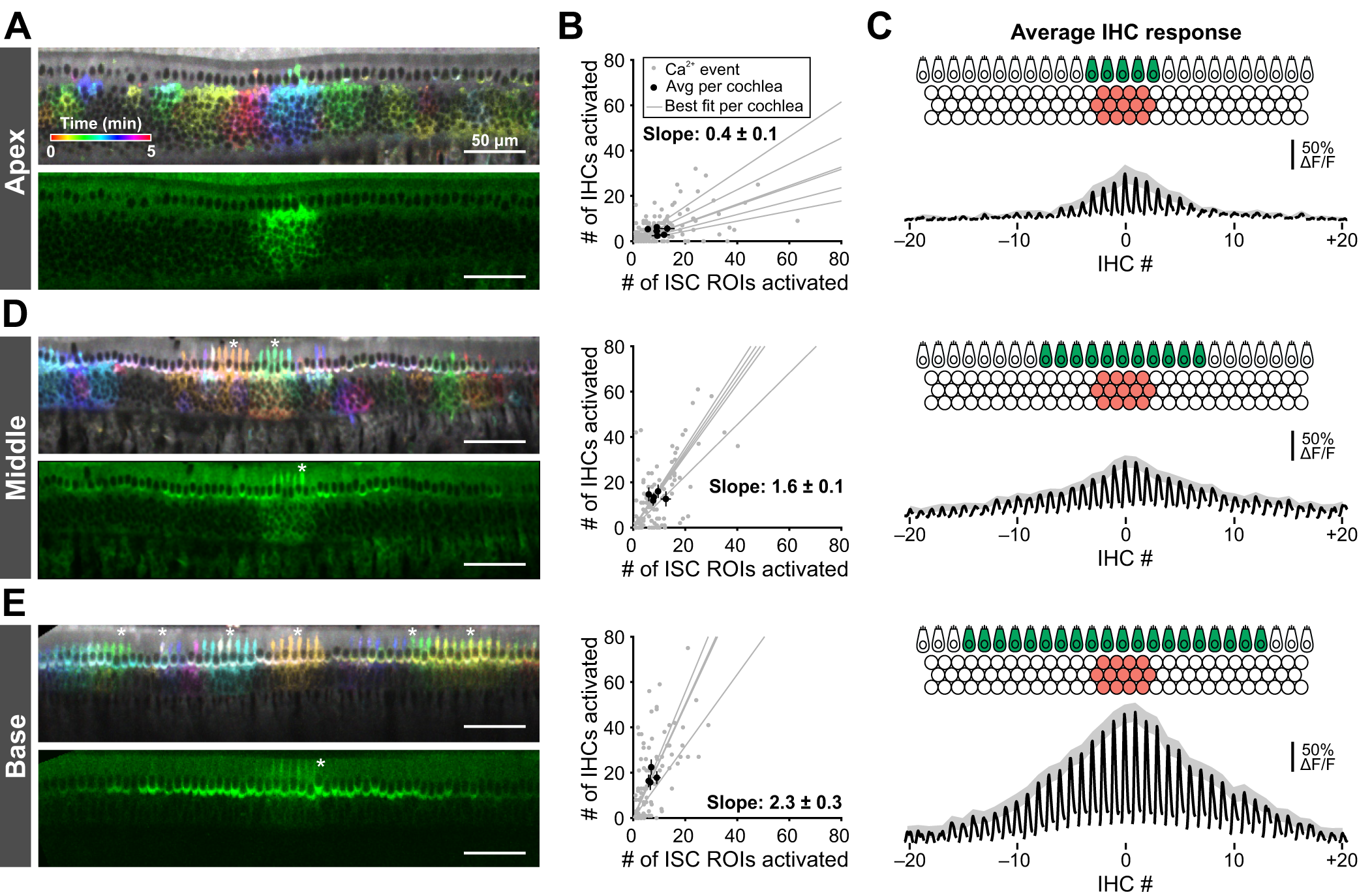
Tonotopic differences in extent of IHC activation at early developmental time points. (A) Images of Ca^2+^ transients colored based on time of occurrence (top) and exemplar Ca^2+^ transient (bottom) in the apical portions of cochleae isolated from P0 *Pax2-Cre;R26-lsl-GCaMP3* mice. Imaging was performed at near physiological temperature (32-34°C). (B) Plot of number of IHCs activated as a function of the number of ISC ROIs activated for apical (top, n = 6 cochleae) regions of the cochlea. Grey dots indicate individual Ca^2+^ transients, grey lines indicated linear best fits for each cochlea, and black dots and lines indicate the mean event size ± SEM for each cochlea. Calculated slope is the mean ± SEM of the best fit lines. (C) Schematic and average IHC response for aligned ISC events in the apical (top), middle (middle), and basal (bottom) regions of the cochlea. Black traces are the average IHC responses to an ISC event with centroid closest to center IHC (IHC at 0). Grey shaded region indicated SEM for the peak of each IHC. (D-E) Similar to A-C, but for middle (n = 5 cochleae), and basal (n = 4 cochleae) portions of the cochlea.

**Figure 7.**
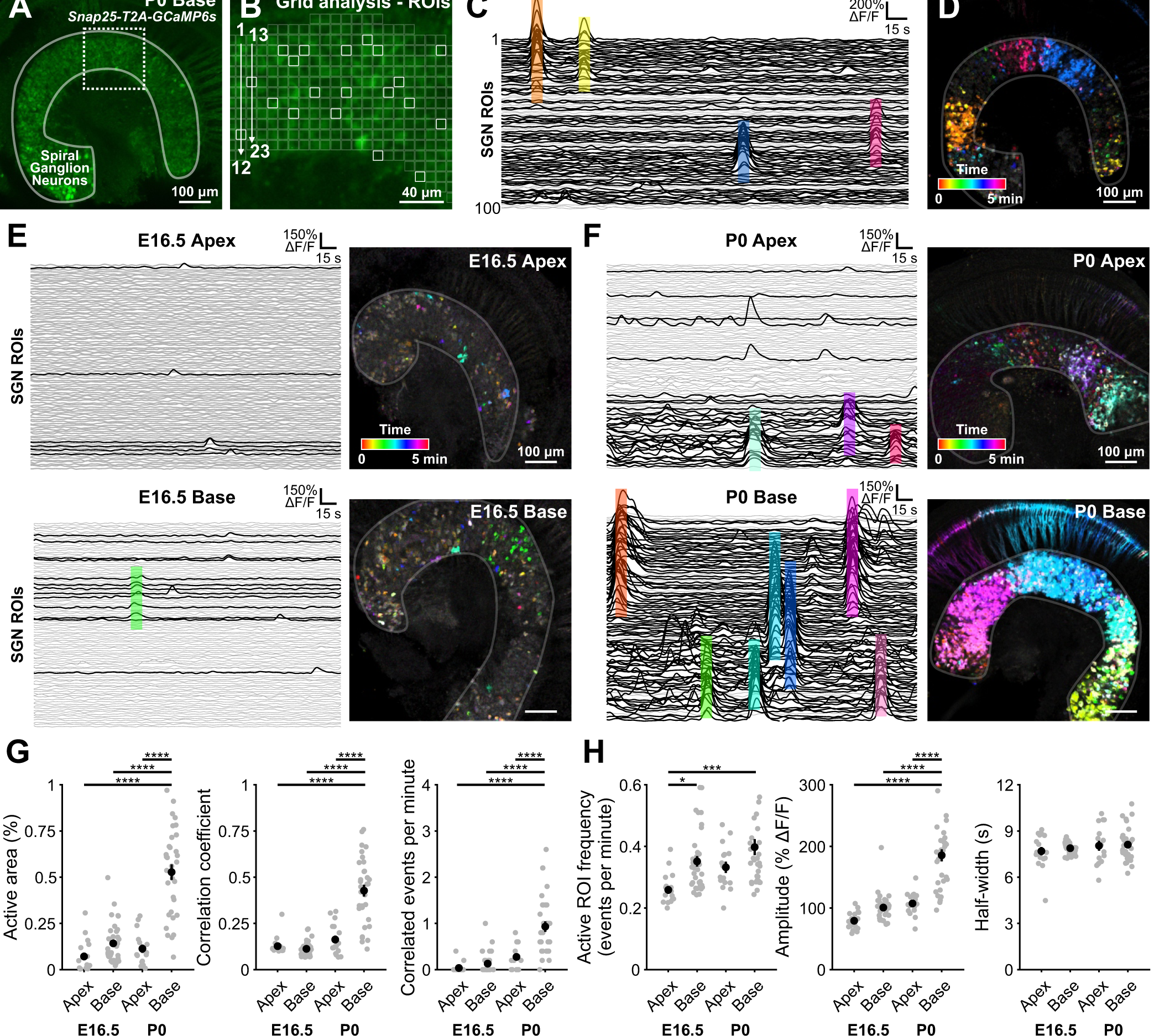
Tonotopic differences in extent of SGN activation at early developmental time points. (A) Image of an excised basal portion of cochlea from a P0 Snap25-T2A-GCaMP6s mouse, which expresses GCaMP6s in SGNs. Dotted line indicates region shown in (B). (B) For analysis of time lapse imaging, a grid of square ROIs was placed over SGNs. ROIs were numbered top-to-bottom, then left-to-right. All ROIs were analyzed, but only random ROIs were chosen to display in figures (white squares). Imaging was performed at room temperature (∼25°C). (C) Individual ROI traces for SGNs (100 randomly selected). Colored boxes are examples of SGN coordinated activity that align with time-color representation in D. Black traces indicate ROIs with at least one detected peak (5^th^ percentile value ± 5 SDs). Grey traces indicate ROIs with no detected peaks. (D) SGN Ca^2+^ transients colored based on time of occurrence. (E) Individual ROI traces and time-color representation of Ca^2+^ transients in E16.5 apical (top) and basal (bottom) regions of the cochlea. Black traces indicate ROIs with at least one detected peak (5^th^ percentile value ± 5 SDs). Grey traces indicate ROIs with no detected peaks. (F) Similar to (E), but in P0 apical (top) and basal (bottom) regions of the cochlea. (G) Quantification of active area (percentage of ROIs with at least one detected peak), correlation coefficient (80^th^ percentile), and correlated events per minute in E16.5 and P0 cochleae. n = 17 E16.5 apical portions from 9 mice, n = 34 E16.5 basal portions from 17 mice, n = 16 P0 apical portions from 8 mice, and n =32 P0 basal portions from 16 mice, (one-way ANOVA with Tukey post-hoc; ****p < 5e-5) (H) Quantification of frequency, amplitude, and duration of Ca^2+^ transients calculated from individual active ROIs. n values are reported in (G), (one-way ANOVA with Tukey post-hoc; ****p < 5e-5,***p < 5e-4,*p < 0.05, all other comparisons not indicated are not significant).

**Figure 8.**
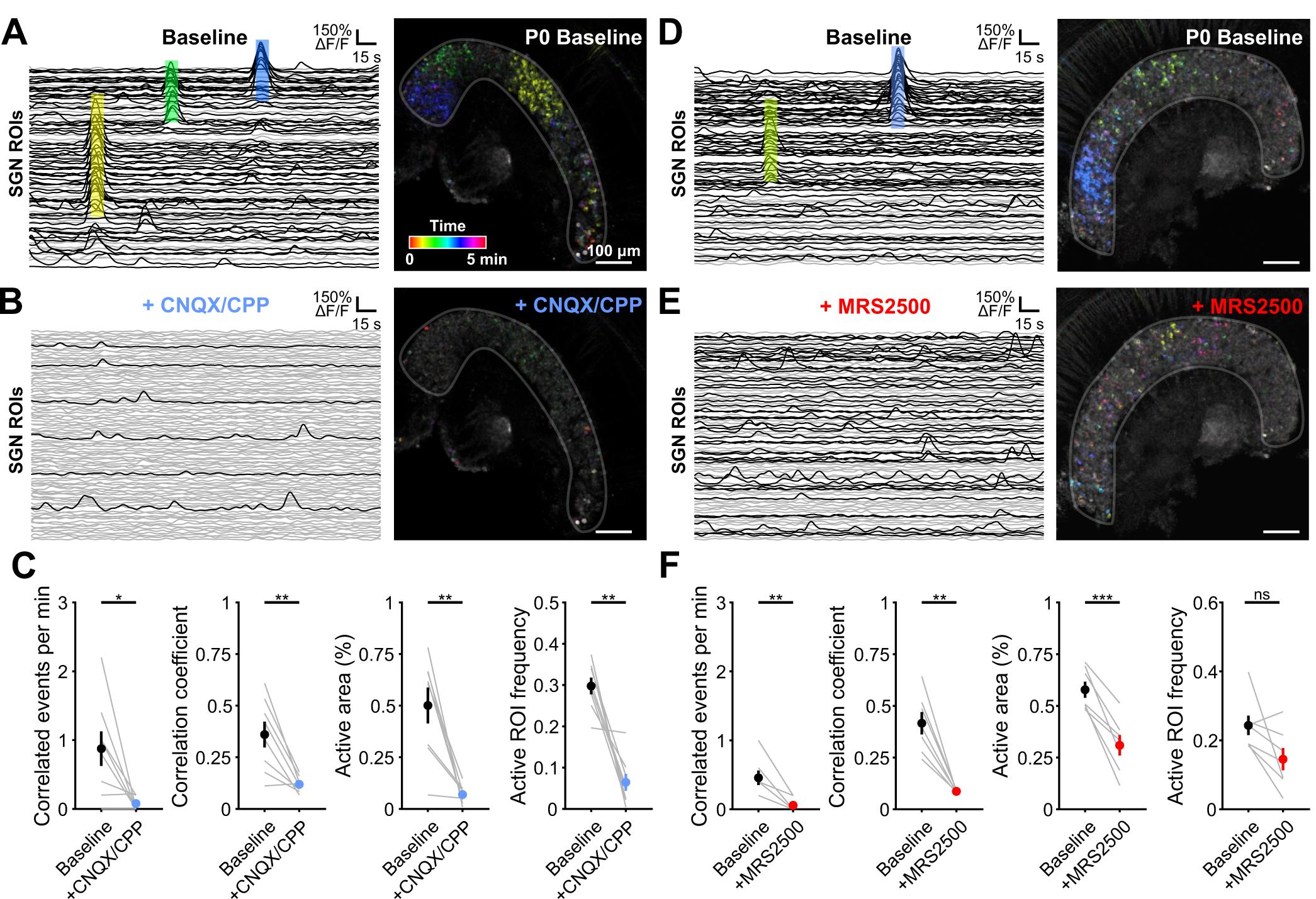
Correlated activation of SGNs requires P2ry1-mediated excitation of IHCs. (A) Individual ROI traces and time-color representation of Ca^2+^ transients in control (baseline) conditions from P0 basal portion of excised cochlea. Colored boxes on left correspond to same colored events on right. Black traces indicate ROIs with at least one detected peak (5^th^ percentile value ± 5 SDs). Grey traces indicate ROIs with no detected peaks. (B) Similar to A, but with application of the AMPAR and NMDAR antagonists CNQX (50 µM) and CPP (100 µM). (C) Quantification of correlated event frequency, correlation coefficient (80^th^ percentile), active area (percentage of ROIs with at least one detected peak), and frequency of transients in active ROIs before and after application of CNQX/CPP. n = 8 P0 basal portions from 4 mice, (paired t-test with Benjamini-Hochberg adjustment; **p < 0.005, *p < 0.05). (D) Similar to A. (E) Similar to D, but with application of the P2RY1 antagonist (MRS2500, 1 μM). (F) Quantification of correlated event frequency, correlation coefficient (80^th^ percentile), active area (percentage of ROIs with at least one detected peak), and frequency of transients in active ROIs before and application of MRS2500. n = 7 P0 basal portions from 4 mice, (paired t-test with Benjamini-Hochberg adjustment; ***p < 5e-4, **p < 0.005, *p < 0.05, ns: not significant).

### Electrophysiology

For inner supporting cell recordings, apical and basal segments of the cochlea were acutely isolated from mouse pups and used within 2 hours of the dissection. Cochleae were moved into a recording chamber and continuously superfused with bicarbonate-buffered artificial cerebrospinal fluid (1.5–2mL/min) consisting of the following (in mM): 119 NaCl, 2.5 KCl, 1.3 MgCl_2_, 1.3 CaCl_2_, 1 NaH_2_PO_4_, 26.2 NaHCO_3_, 11 D-glucose and saturated with 95% O_2_ / 5% CO_2_ to maintain a pH of 7.4. Near physiological temperature (32-34°C) solutions were superfused using a feedback-controlled in-line heater (Warner Instruments). Whole-cell recordings of inner supporting cells (ISCs) were made under visual control using differential interference contrast microscopy (DIC). Electrodes had tip resistances between 3.5-4.5 MΩ when filled with internal solution consisting of (in mM): 134 KCH_3_SO_3_, 20 HEPES, 10 EGTA, 1 MgCl_2_, 0.2 Na-GTP, pH 7.3. Spontaneous currents were recorded with ISCs held at −80 mV and recorded for at least 5 minutes with pClamp10 software using a Multiclamp 700B amplifier, low pass filtered at 1kHz, and digitized at 5kHz with a Digidata 1322A analog-to-digital converter (Axon Instruments). Errors due to voltage drop across the series resistance and the liquid junction potential were left uncompensated for recordings of spontaneous activity. For quantification of spontaneous events, traces were imported into MATLAB and baseline corrected using the msbackadj function (30 s window). Events were defined as peaks in the signal that exceed 20 pA using the findpeaks function (minPeakProminence = 20). Amplitude is represented as the mean amplitude and integral as the average charge transfer per second (pA/s). Input resistances were calculated by taking the change in voltage to a small negative current injection and dividing it by the amplitude of the current injection (–100 pA). For experiments with MRS2500 application, a 5-minute baseline was collected before beginning flow of MRS2500 (1 uM). After a 3-minute wash in period, the following 5-minute period was used for MRS2500 analysis.

### Immunohistochemistry

Mice were deeply anesthetized with isoflurane and perfused with freshly prepared paraformaldehyde (4%) in 0.1 M phosphate buffer. Cochleae were post-fixed overnight at room temperature and stored at 4°C in PBS until processing. For immunohistochemistry, P0-P11 cochleae were removed from the temporal bone and washed 3 x 5 minutes with PBS. Cochleae were incubated overnight with primary antibodies against β-gal (anti-Rabbit; 1:4000, Sanes laboratory) and calbindin (anti-goat; 1:500, Santa Cruz) for detection of β-gal and visualization of hair cells. Cochleae were then rinsed three times with PBS and incubated for two hours at room temperature with secondary antibodies raised in donkey (Alexa-488 and Alexa-546; 1:2000, Life Technologies). Slides were washed three times in PBS (second with PBS + 1:10,000 DAPI), allowed to dry, and sealed using Aqua Polymount (Polysciences, Inc.). Images were captured using a laser scanning confocal microscope (LSM 880, Zeiss).

### Transmitted Light Imaging

Cochlear segments were imaged with an Olympus 40x water immersion objective (LUMPlanFl/IR) and recorded using MATLAB and a USB capture card (EZ Cap). Movies were generated by subtracting frames at time t_n_ and t_n+5_ seconds using MATLAB. To quantify transmittance changes, a threshold of three standard deviations above the mean was applied to each pixel value over time. To calculate the frequency of these events, the whole field was taken as an ROI and peaks detected using MATLAB (findpeaks function). For experiments with MRS2500 application, a 10-minute baseline was collected before beginning flow of MRS2500 (1 uM). After a 5-minute wash in period, the following 10-minute period was used for MRS2500 analysis. An additional washout period of 25 minutes was captured, with the final 10 minutes quantified as washout.

### Cochlear explant culture

For imaging of ISCs and IHCs, cochleae were dissected from postnatal day 0 *Pax2-Cre;R26-lsl-GCaMP3* mice in ice-cold, sterile-filtered HEPES-buffered artificial cerebrospinal fluid (aCSF) consisting of the following (in mM): 130 NaCl, 2.5 KCl, 10 HEPES, 1 NaH_2_PO_4_, 1.3 MgCl_2,_ 2.5 CaCl_2_, and 11 D-Glucose. Explants were mounted onto Cell-Tak (Corning) treated coverslips and incubated at 37°C for 2-6 hours in Dulbecco’s modified Eagle’s medium (F-12/DMEM; Invitrogen) supplemented with 1% fetal bovine serum (FBS) and 10U/mL penicillin (Sigma) prior to imaging.

For imaging of SGNs, a detailed cochlear culture protocol has been described previously (Driver and Kelley, 2010). Briefly, cochleae collected from *Snap25-T2A-GCaMP6s* E16.5 and P0 embryos were dissected in 1X Hank’s buffered saline solution (HBSS)/HEPES and separated into apical and basal pieces. Cochlear pieces were transferred onto polycarbonate membrane filters (Sterlitech PCT0213100) in a 14mm bottom well dish with #0 cover glass (In Vitro Scientific, D29-14-0-N) filled with 250 µL media containing L-15 media (Invitrogen, 21083027), 10% fetal bovine serum, 0.2% N2, 0.001% ciprofloxacin, and 0.1 mM Trolox. Cochlear pieces were flattened by surface tension and incubated at 37 degree with 95% O_2_/5% CO_2_ for a minimum of 2 and maximum of 6 hours before imaging.

### Confocal imaging of explants

For imaging of ISCs and IHCs, cochleae were moved into a recording chamber and continuously superfused with bicarbonate-buffered artificial cerebrospinal fluid (1.5 - 2 mL/min) consisting of the following (in mM): 115 NaCl, 6 KCl, 1.3 MgCl_2_, 1.3 CaCl_2_, 1 NaH_2_PO_4_, 26.2 NaHCO_3_, 11 D-glucose, and saturated with 95% O_2_ / 5% CO_2_ to maintain a pH of 7.4. Images were captured at 2 frames per second using a Zeiss laser scanning confocal microscope (LSM 710, Zeiss) through a 20X objective (Plan APOCHROMAT 20x/1.0 NA) at 512 x 512 pixels (354 x 354 µm; 16-bit depth) resolution. Sections were illuminated with a 488 nm laser (maximum 25 mW power). MRS2500 (1 µM, Tocris) was applied by addition to superfused ACSF.

For imaging of SGNs, cultured cochlear pieces were removed from the incubator and placed with SGNs/IHCs facing down. Cochlear pieces were stabilized with a platinum harp with nylon strings. The bottom of the well was filled with 250 µL static bath of artificial cerebrospinal fluid (ACSF) consisting of the following (in mM): 145 NaCl, 5 KCl, 1 CaCl2, 1 MgCl2, 1 NaH2PO4, 5 HEPES, and 5 D-glucose with a pH of 7.4. Imaging was performed at room temperature (22 – 24°C). Images were captured at 1 frame per second using a Zeiss laser scanning confocal microscope (LSM 880, Zeiss) through a 20X objective (Plan-Apochromat 20x/0.8 M27) at 800 x 800 pixels (708 µm x 708 µm; 16-bit depth, 1.32 µs dwell time) resolution. Tissues were illuminated with a 488 nm Argon laser with emission ranging 500-540 nm and a GaAsp detector. Baseline imaging sessions consisted of 5 consecutive minutes of recording. Image acquisition was stopped and CNQX (50 µM; Sigma, C239) and CPP (100 µM; Abcam, ab120159) or MRS2500 (1 µM; Tocris, 2154) were added directly to the bath and allowed to equilibrate for 5 minutes before capturing an additional 5-minutes used for analysis of pharmacological block.

### Analysis of in vitro Ca^2+^ transients

For analysis of IHC and ISC activity, image stacks were imported into MATLAB where a region of interest was drawn around the ISC and IHCs. For ISCs, a 10 pixel by 10 pixel grid was imposed across the entire image and only squares within the drawn ISC ROI region were analyzed. The signal-to-noise ratio was extremely high within individual ISC ROIs (Figure 5E) and ISC events were defined by contiguous activation of connected ISC ROIs (all 26 edges and vertices of each timepoint per ROI were considered; see Supplemental Video 2). IHCs were semi-automatically detected by finding local minima within the IHC ROI and validated by the experimenters. Individual circular ROIs were drawn at the basal pole of each IHC. Fluorescence changes were normalized as ΔF/F_o_ values, where ΔF = F - F_o_ and F_o_ was defined as the fifth percentile value for each pixel. Peaks in the signals were detected in MATLAB using the built-in peak detection function (findpeaks) with a fixed value threshold criterion (median + 3 SDs for ISCs and median + 4 SDs for IHCs). IHC activity was considered coincident if ISC and IHC were co-active in both space and time. IHC coordinated events were defined as anytime more than 3 adjacent IHCs were co-active at the same time. Correlation coefficients reported are the 80th percentile correlation coefficient. For analysis of the extent of IHC activation, ISC events were centered around the center of mass for each event. The IHC closest to the center of mass was designated as IHC 0 and the adjacent 20 IHCs on either side were examined. If the adjacent 20 IHCs included IHCs that were not within the imaging frame (i.e. if ISC event occurred near the edge), visible IHCs were averaged while out-of-frame IHCs were not.

For analysis of SGN signals, image stacks were imported into MATLAB where a region of interest was drawn around the SGNs. A 10 pixel by 10 pixel grid was imposed across the entire image and only squares within the drawn SGN ROI region were analyzed. Fluorescence changes were normalized as ΔF/F_o_ values, where ΔF = F - F_o_ and F_o_ was defined as the fifth percentile value for each pixel. Peaks in the signals were detected in MATLAB using the built-in peak detection function (findpeaks) with a fixed value threshold criterion (5^th^ percentile value + 5 SDs). Active area is defined as the percentage of active ROIs (ROIs with at least 1 peak) within the drawn SGN region. Correlation coefficient was defined as the 80^th^ percentile correlation coefficient among active ROIs only. Correlated events were defined as coincident SGN activation of 35 SGN ROIs. While this parameter (35 SGNs for a correlated event) was subjective, re-analysis of the data where this parameter was varied (down to 15 ROIs for a correlated event) revealed that relative frequencies of correlated events were preserved between conditions. Active ROI frequency, amplitude, and half-widths were calculated using only active ROIs.

### Installation of cranial windows

Inhalation anesthesia was induced with vaporized isoflurane (4% for 5 minutes, or until mice are non-responsive to toe-pinch) and surgical plane maintained during the procedure (with 1-2% isoflurane) with a stable respiration rate of 80 breaths per minute. A midline incision beginning posterior to the ears and ending just anterior to the eyes was made. Two subsequent cuts were made to remove the dorsal surface of the scalp. A headbar was secured to the head using super glue (Krazy Glue). Fascia and neck muscles overlying the interparietal bone were resected and the area bathed in sterile, HEPES-buffered artificial cerebrospinal fluid that was replaced as necessary throughout the surgery. Using a 28G needle and microblade, the sutures circumscribing the interparietal bone were cut and removed to expose the midbrain. The dura mater was removed using fine scissors and forceps, exposing the colliculi and extensive vasculature. A 5 mm coverslip (CS-5R; Warner Instruments) was then placed over the craniotomy, the surrounding bone was dried using a Kimwipe, and super glue was placed along the outer edges of the coverslip for adhesion to the skull. Replacement 0.9% NaCl solution was injected IP and a local injection of lidocaine was given to the back of the neck. Animals were weaned off isoflurane, placed under a warming lamp, and allowed to recover for a minimum of 1 hour prior to imaging.

### In vivo calcium imaging

After 1 hour of post-surgical recovery from anesthesia, pups were moved into a swaddling 15 mL conical centrifuge tube. The top half of this tube was removed to allow access to the headbar and visualization of the midbrain or midbrain and caudal part of the cortex. Pups were head-fixed and maintained at 37°C using a heating pad and temperature controller (TC-1000; CWE). During the experiments, pups were generally immobile; however, occasional limb and tail twitching did occur.

For wide field epifluorescence imaging, images were captured at 10 Hz using a Hamamatsu ORCA-Flash4.0 LT digital CMOS camera attached to a Zeiss Axio Zoom.V16 stereo zoom microscope. A 4 x 4 mm field of view was illuminated continuously with a mercury lamp (Zeiss Illuminator HXP 200C) and visualized through a 1X PlanNeoFluar Z 1.0x objective at 17x zoom. Images were captured at a resolution of 512 x 512 pixels (16-bit pixel depth) after 2 x 2 binning to increase sensitivity. Each recording consisted of uninterrupted acquisition over 30 minutes or 40 minutes if injected with pharmacological agents.

### Catheterization of animals for in vivo imaging

After induction of anesthesia and before installing the cranial window, a catheter was placed in the intraperitoneal (IP) space of neonatal mouse pups. A 24G needle was used to puncture the peritoneum and a small-diameter catheter (SAI Infusion Technologies, MIT-01) was placed. A small drop of Vetbond secured the catheter to the pup’s belly. Installation of cranial window proceeded as described above.

Imaging sessions consisted of 15 minutes of baseline activity measurements, followed by a slow push of either 50 µL of sham (5% mannitol solution) or MRS2500 solution (500 µM in 5% mannitol solution). Imaging was continuous throughout and 45 minutes of activity total were collected. No discernable diminishment of activity was observed in sham animals.

### Image processing

For wide field imaging, raw images were imported into the MATLAB environment and corrected for photobleaching by fitting a single exponential to the fluorescence decay and subtracting this component from the signal. Intensities were normalized as ΔF/F_o_ values, where ΔF = F - F_o_ and F_o_ was defined as the fifth percentile value for each pixel. Ovoid regions of interest (ROIs) encompassing the entire left and right inferior colliculi were drawn. Across all conditions, the size of the ROIs was invariant. However, due to small differences in the imaging field between animals, the ROIs were placed manually for each imaging session. Peaks in the signals were detected in MATLAB using the built-in peak detection function (findpeaks) using a fixed value threshold criterion; because fluorescence values were normalized, this threshold was fixed across conditions (2% ΔF/F_o_). Occasionally, large events in the cortex or superior colliculus would result in detectable fluorescence increases in the IC. These events broadly activated the entire surface of the IC and did not exhibit the same spatially-confined characteristics as events driven by the periphery. These events were not included in the analysis.

### Analysis of retinal wave activity in the superior colliculus

ROIs (200 x 150 pixels) were placed over each lobe of the superior colliculus and downsampled by a factor of five. Signals were normalized as ΔF/F_o_ values, where ΔF = F - F_o_ and F_o_ was defined as the fifth percentile value for each pixel. In order to eliminate periodic whole-sample increases in fluorescence, the mean intensity of all pixels was subtracted from each individual pixel. Following this, pixels were considered active if they exceeded the mean + 3 standard deviations. For each point in time, the number of active pixels was summed. Retinal waves were defined as prolonged periods (> 1 second), where more than 5 pixels were active simultaneously. Retinal wave durations were defined as the total continuous amount of time that more than 5 pixels were active.

### Experimental design and statistical analysis

All statistics were performed in the MATLAB (Mathworks) programming environment. All statistical details, including the exact value of n, what n represents, and which statistical test was performed, can be found in the figure legends. To achieve statistical power of 0.8 with a 30% effect size with means and standard deviations similar to those observed in previous studies (Figure 1E of Tritsch et al., 2007 and Figure 1B, 3D in Wang et al., 2015), power calculations indicated that 7 animals in each condition were necessary (µ_1_ = 10, µ_2_ = 7, σ = 2, sampling ratio = 1). While this number was used as a guide, power calculations were not explicitly performed before each experiment; many experiments had much larger effect sizes and sample sizes were adjusted accordingly. For transparency, all individual data points are included in the figures. Data are presented as mean ± standard error of the mean (SEM). Because the main comparison between conditions was the mean, the SEM is displayed to highlight the dispersion of sample means around the population mean. All datasets were tested for Gaussian normality using the D’Agostino’s K^2^ test. For single comparisons, significance was defined as p <= 0.05. When multiple comparisons were made, the Benjamini-Hochberg or Bonferroni correction was used to adjust p-values accordingly to lower the probability of type I errors. For multiple condition datasets, one-way ANOVAs were used, followed by Tukey’s multiple comparison tests.

## RESULTS

### Spontaneous electrical activity of inner supporting cells emerges before birth

In the developing mammalian cochlea, supporting cells within Kölliker’s organ spontaneously release ATP, initiating a purinergic signaling cascade that releases Ca^2+^ from intracellular stores and activates TMEM16A, a Ca^2+^-activated Cl^−^ channel (Wang et al., 2015; Babola et al., 2020). Efflux of Cl^−^ ions draws positive K^+^ ions into the extracellular space, producing a temporary osmotic gradient that draws water into the extracellular space. Because of extensive gap-junction coupling between ISCs, activation of these purinergic pathways induces large currents and cellular shrinkage (crenation) among groups of these cells. Local increases in extracellular K^+^ depolarize nearby IHCs, resulting in bursts of action potentials, glutamate release, and activation of AMPA and NMDA receptors on post-synaptic SGNs (Zhang-Hooks et al., 2016); thus, unlike hearing, this electrical activity does not require activation of mechanotransduction channels (Sun et al., 2018). Spontaneous inward currents in ISCs are present from birth, but little is known about when this activity emerges (Tritsch and Bergles, 2010; Wang et al., 2015; Zhang-Hooks et al., 2016). To determine the onset of spontaneous ISC currents, we made whole-cell voltage clamp recordings from inner supporting cells (ISCs) in cochleae acutely isolated from embryonic day 14 to 16 (E14-16) mouse pups (Figure 1A), a developmental period characterized by basal to apical differentiation of inner and outer hair cells (Chen et al., 2002). Recordings from ISCs in the apical region of the cochleae revealed no discernable spontaneous currents (6/6 cochleae; Figure 1B). In contrast, large spontaneous currents were observed in most cochleae in the basal region (3/3 cochleae at E16 and 1/3 cochleae at E14). After birth, spontaneous inward currents were observed in apical and basal ISCs throughout the early postnatal period (Figure 1C), consistent with previous observations (Tritsch et al., 2007, Tritsch et al., 2010). At P0, these currents were more frequent (24 ± 2 versus 1 ± 1 events per minute; One-way ANOVA, F(3,32) = 27.11, p = 6e-9; Tukey HSD, p = 6e-9), larger in amplitude (522 ± 100 versus 44 ± 10 pA; One-way ANOVA, F(3,32) = 4.14, p = 0.013; Tukey HSD, p = 0.007), and carried more charge per second (integral; 320 ± 80 versus 15 ± 5 pC; One-way ANOVA, F(3,32) = 0.036; Tukey HSD, p = 0.022) than in embryonic ISCs (Figure 1D). While there was a progressive decline in average frequency, amplitude, and integral postnatally up to hearing onset (∼P12), these changes were not statistically significant (Figure 1D). The lack of spontaneous currents at embryonic ages may reflect that ISCs are not as extensively coupled by gap junctions, which would prevent detection of currents that arise in distant cells. However, the membrane resistances of apical ISCs were consistently low (Figure 1D; E14-16: 11 ± 2 MΩ, P0-2: 11 ± 4 MΩ, P7-8: 9 ± 3 MΩ, P10-12: 14 ± 4 MΩ; One-way ANOVA, F(3,32) = 0.38, p = 0.77). As membrane resistance is determined primarily by cell to cell coupling (Jagger and Forge, 2006), these results suggest that gap junctional coupling among ISCs across this developmental period is similar (Jagger and Forge, 2006; Kamiya et al., 2014). Together, these data indicate that spontaneous currents emerge in ISCs during the late embryonic period in a basal to apical gradient, matching the progression of hair cell maturation.

### Supporting cell spontaneous currents and crenation are mediated by P2RY1 throughout the prehearing period

Spontaneous currents in ISCs require activation of purinergic receptors between birth and shortly after hearing onset, when Kölliker’s organ recedes (Tritsch and Bergles, 2010). Recently, the G_q_-coupled P2Y1 receptor (P2RY1) was identified as the primary purinergic autoreceptor mediating spontaneous currents in ISCs after the first postnatal week. Gene expression studies revealed that *P2ry1* is expressed at much higher levels (>100 fold) than any other P2Y receptor in the cochleae, even at early embryonic ages (Scheffer et al., 2015; Kolla et al., 2020), suggesting that this receptor may initiate spontaneous currents throughout development. However, the presence of Ca^2+^-permeable ionotropic (P2×2/4) and G_q_-coupled metabotropic receptors (P2RY2/4/6) in the cochlea (Huang et al., 2010; Housley et al., 2013; Kolla et al., 2020) indicate that alternative pathways could also contribute to spontaneous activity during this period, depending on the amount, location and kinetics of ATP release, as well as the presence and activity of extracellular nucleotidases. To define the dynamics of P2RY1 expression during cochlear development, we isolated cochleae from *P2ry1-LacZ* reporter mice and performed immunostaining for β-galactosidase at different developmental ages (Figure 2). Immunofluorescence within Kölliker’s organ and along the entire length of the cochlea was detected across all postnatal time points (P0, P7 and P11). At later stages of development, β-galactosidase immunofluorescence was observed in interdigitating phalangeal cells, primarily within the base at P7 and by P11 along the entire length of the cochlea (Figure 2). These data indicate that P2RY1 promoter activity is localized to ISCs throughout the prehearing period, providing the means to express P2RY1 and detect ATP release from these cells.

To determine if P2RY1 mediates spontaneous currents in ISCs across this developmental period, we examined the sensitivity of these responses to the specific P2RY1 antagonist, MRS2500 (Houston et al., 2006), which displays no obvious off-target effects in cochleae isolated from P2RY1 knockout mice (Babola et al., 2020). At the earliest time points exhibiting robust spontaneous activity (E16-17), acute inhibition of P2RY1 with MRS2500 (1 µM) dramatically reduced the frequency (baseline: 15 ± 3, MRS2500: 3 ± 1 events per minute; Student’s t-test with Bonferroni correction, t(6) = 5.36, p = 0.0017), amplitude (baseline: 280 ± 40, MRS2500: 94 ± 20 pA; Student’s t-test with Bonferroni correction, t(6) = 4.37, p = 0.0047), and total charge transfer of spontaneous inward currents (baseline: 110 ± 30, MRS2500: 20 ± 4 pC; Student’s t-test with Bonferroni correction, t(6) = 3.43, p= 0.0140) (Figure 3A,B). Spontaneous currents were also largely inhibited by MRS2500 at P0, P7, and just prior to hearing onset (P10-12); only small amplitude currents persisted in the presence of this antagonist, consistent with previous observations of residual non-purinergic mediated currents in these cells (Babola et al, 2020).

The efflux of K^+^ and Cl^−^ following purinergic receptor activation induces osmotic shrinkage (crenation) of ISCs through movement of water down its osmotic gradient (Tritsch et al., 2007; Wang et al., 2015). Previous studies revealed that crenations are small and infrequent at birth, but rapidly increase in frequency and size over the first postnatal week (Tritsch, 2010). To determine if P2RY1 mediates cellular crenation throughout development, we monitored acutely isolated apical portions of the cochlea using differential contrast imaging (DIC) before and after application of the P2RY1 antagonist, MRS2500. Crenations were absent in embryonic and P0 mouse pups (Figure 4A) and application of MRS2500 at these ages resulted in no change in the optical properties of the tissue (Figure 4B, Supplemental Video 1). In contrast, crenations were robust at P7 and P11, with the majority of events occurring near the medial edge of IHCs (Figure 4B). Crenation at these ages was reversibly blocked by MRS2500 (Figure 4B, Supplemental Video 1). Together, these results indicate that P2RY1 mediates ISC spontaneous currents throughout the prehearing period and ISC crenation when they emerge after the first post-natal week.

### Correlated activation of IHCs requires activation of ISC P2Y1 receptors

The rapid increase in extracellular K^+^ following activation of ISC purinergic autoreceptors depolarizes nearby IHCs, resulting in high frequency burst firing that triggers glutamate release and subsequent activation of SGNs. Previous studies revealed that activation of P2RY1 autoreceptors is required to induce coordinated activation of groups of ISCs and nearby IHCs after the first postnatal week (Babola et al., 2020). To evaluate if P2RY1 initiates coordinated activity patterns in ISCs and IHCs at earlier developmental time points, we monitored large-scale activity patterns in excised cochleae from P0 *Pax2-Cre;R26-lsl-GCaMP3* mice, which express GCaMP3 in nearly all cells of the cochlea (Figure 5A). We quantified activity patterns by placing a grid composed of square regions of interest (10 x 10 pixels) over the ISC region and circular regions of interest (ROIs) around the basal pole of each IHC, where Ca_v_1.3 Ca^2+^ channels enable depolarization-induced Ca^2+^ influx (Figure 5A,B) (Brandt et al., 2003; Zampini et al., 2014). Time lapse imaging revealed robust spontaneous Ca^2+^ transients in ISCs and concurrent activation of nearby IHCs (Figure 5C,E and Supplemental Video 2). These coordinated transients were abolished following inhibition of P2RY1 with MRS2500 (Figure 5D,E and Supplemental Video 3). At later postnatal ages, persistent inhibition of P2RY1 results in a gradual increase in non-correlated activity in IHCs, due to an accumulation of extracellular K^+^ (Babola et al., 2020). Consistent with this finding, non-correlated IHC activity also emerged after prolonged P2RY1 inhibition in P0 cochleae (Figure 5D-F). These data indicate that early coordinated activation of ISCs and IHCs also requires activation of P2RY1 signaling pathways.

The early postnatal period is defined by transformation of Kölliker’s organ into the inner sulcus (Hinojosa, 1977) and rapid changes in the electrophysiological properties of IHCs, both of which occur in a basal to apical developmental gradient. To determine how these processes affect the activity patterns of IHCs, we assessed IHC activation in apical, middle, and basal portions of cochleae from P0 *Pax2-Cre;R26-lsl-GCaMP3* mice. In the apex, groups of ISCs exhibited robust coordinated Ca^2+^ transients that occurred along the entire length and medial-lateral portion of the imaged area (Figure 6A). For each individual ISC event, only IHCs within the immediate area were activated (4.5 ± 0.6 IHCs per ISC event; Figure 6A,B and Supplemental Video 4). To determine how the area of ISC activation influences the number of hair cell activated, we examined the relationship between the number of ISC ROIs activated and the number of IHCs activated (Figure 6B). The relationship was linear, with more IHCs active following large ISC events; however, ISCs did not strongly activate IHCs in the apex (0.4 ± 0.1 IHCs activated per single ISC ROI; Figure 6B). We then computationally centered each ISC event to explore how IHC activation varies as a function of distance away from the center of each ISC event (Figure 6C). On average, IHCs in the apex were moderately activated following ISC activation, with few IHCs activated distal to the event. In contrast, in the developmentally older middle and basal portions of the cochlea, progressively more IHCs were activated on average for each ISC event (13.8 ± 0.7 and 18.1 ± 1.5 IHCs per ISC event, respectively; Figure 6D,E). Each IHC exhibited larger Ca^2+^ transients (base: 310 ± 10% ΔF/F for center IHC, middle: 160 ± 10% ΔF/F, and apex: 100 ± 20% ΔF/F; One-way ANOVA, F(2,172) = 80.95, p = 2e-25), and IHC activation extended far beyond the active ISCs region (Figure 6D,E and Supplemental Video 4). We did not observe any difference between the average number of ISCs activated per event (base: 7.1 ± 0.6 ISCs, middle: 8.7 ± 1.1 ISCs, apex: 9.6 ± 1.0 ISCs; One-way ANOVA, F(2,12) = 1.45, p = 0.27) or the average event amplitude (base: 82 ± 5 % ΔF/F, middle: 106 ± 8 % ΔF/F, apex: 101 ± 6 % ΔF/F; One-way ANOVA, F(2,12) = 2.6, p = 0.11), suggesting that changes in ISC activity are not responsible for the difference in IHC activation along the tonotopic axis. However, phalangeal cells, which envelop IHCs, displayed prominent Ca^2+^ transients coincident with those in Kölliker’s organ in the basal and middle turns (asterisks in Figure 6D,E), but not in the apical turn (Figure 6A), which may contribute to the muted response of apical IHCs. Together, these data indicate that IHCs in basal portions of the cochlea are activated by ISCs at an earlier developmental stage.

### Correlated activation of SGNs requires P2RY1-mediated excitation of IHCs

Previous studies indicate that burst firing of SGNs during the prehearing period requires glutamatergic synaptic excitation (Seal et al., 2008). Within apical portions of the cochlea, SGN afferent fibers extend into the newly differentiated hair cell region at E16, but SGNs do not exhibit post-synaptic densities and IHCs do not form ribbons until E18 (Michanski et al., 2019), suggesting that IHC activity may not propagate to the CNS at this stage. To determine when P2RY1-mediated currents in ISCs trigger coordinated activation of SGNs, we performed time-lapse imaging of excised cochleae from mice that expressed GCaMP6s in SGNs (*Snap25-T2A-GCaMP6s* mice) (Figure 7A). Similar to the analysis of ISC activity, we placed a grid of square ROIs over SGNs to monitor changes in fluorescence over time across the population (Figure 7B-D). Consistent with the lack of activity in apical ISCs at E16 (Figure 1C), SGN Ca^2+^ transients were infrequent and non-correlated at this age (Figure 7E). In the base, where ISCs exhibit robust ATP-mediated currents (Figure 1C), SGNs were also largely silent, with some preparations (10/32) exhibiting infrequent, concurrent activation of groups of SGNs (Figure 7E). However, at P0, most SGNs at the basal end of apical preparations exhibited correlated activation (10/16 preparations, Figure 7F), consistent with the base-to-apex emergence of activity in ISCs. Compared to E16 and P0 apical preparations, P0 basal preparations had larger average numbers of SGNs activated, higher correlations among ROIs, and more frequent correlated events (Figure 7G and Supplemental Video 5), although no differences were observed in the duration of events (Figure 7H). Considering only active ROIs from each preparation, transients from P0 basal preparations were more frequent than E16.5 apical preparations and were larger in amplitude than all other preparations (Figure 7H). These data indicate that coordinated activation of SGNs emerges between E16.5 and P0 in a basal to apical developmental gradient.

Extrusion of K^+^ into the extracellular space following P2RY1 activation non-selectively depolarizes nearby cells and their processes, including SGN dendrites (Tritsch et al., 2007). Although synaptic excitation is required to induce burst firing of SGNs in wild type mice, homeostatic increases in SGN membrane resistance and thereby excitability in deaf mice (*Vglut3* KOs) allows direct activation of groups of SGNs by these brief elevations of extracellular K^+^ (Babola et al., 2018). Given that the membrane resistance of SGNs is extremely high at birth (Marrs and Spirou, 2012), it is possible that extruded K^+^ could directly drive the activity of nearby SGNs at earlier developmental time points. To determine if coordinated SGN Ca^2+^ transients require release of glutamate from IHCs, we applied the AMPA and NMDA receptor antagonists CNQX and CPP to acutely isolated P0 preparations of basal cochleae (Figure 8A,B and Supplemental Video 6). The number of coordinated events, the correlation coefficient between ROIs, the number of active ROIs, and the average frequency of transients per ROI were markedly decreased by CNQX/CPP (Figure 8C), indicating that coordinated activation of SGNs at this early developmental stage also requires activation of ionotropic glutamate receptors. While coordinated SGN transients were abolished, individual SGNs exhibited infrequent Ca^2+^ transients when deprived of glutamatergic excitation, suggesting that there is a form of activity that is independent of synaptic excitation (Figure 8B); however, this activity was not coordinated between neighboring SGNs, suggesting that it may arise through cell intrinsic processes.

Given the dependence of coordinated IHC activation on P2RY1-mediated ISC activity (Figure 5B,C), coordinated SGN activity should also be sensitive to P2RY1 inhibition. Indeed, application of MRS2500 decreased the number of coordinated SGN transients, the correlation coefficient between ROIs, and the number of active ROIs (Figure 8D-F and Supplemental Video 7). The average ROI transient frequency did not decrease in MRS2500, consistent with observations of increased, uncorrelated activity of IHCs with prolonged P2RY1 inhibition (Figure 5E). Together, these data indicate that activation of P2RY1 on ISCs leads to IHC depolarization, glutamate release, and post-synaptic activation of SGNs when functional synapses first emerge at ∼P0.

### Developmental changes in spontaneous activity in the inferior colliculus

In the developing auditory midbrain (inferior colliculus), bursts of activity originating in the cochlea coordinate the activity of neurons within isofrequency lamina (Babola et al., 2018), regions later responsive to specific frequencies of sounds. Events arising within one cochlea induce bilateral activity in both lobes of the IC, with the contralateral lobe exhibiting the strongest response, consistent with the known contralateral bias in information flow through the auditory pathway. While bursts of electrical activity have been detected as early as P1 in the auditory brainstem in anesthetized animals (Tritsch et al., 2010), little is known about how the spatial and temporal aspects of this activity change *in vivo* with development. To define developmental changes in IC activity, we performed time-lapse imaging of awake mice in *Snap25-T2A-GCaMP6s* mice. At all ages examined (from P1, the earliest age we could reliably perform imaging, to P16, just after hearing onset), periodic excitation of neurons occurred within isofrequency domains of both lobes of the IC (Figure 9A,B and Supplemental Video 8). Bilateral events stochastically alternated between having larger amplitudes on the right and left, indicative of electrical activity coming from the left or right cochlea respectively (Babola et al., 2018). The degree of lateralization (smaller/larger amplitude) also varied on an event-by-event basis (degree of left/right dominance represented by dot sizes in Figure 9C). On average, events were evenly balanced between left and right IC (Figure 9C) and increased in frequency and amplitude with developmental age (Figure 9D). The correlation between activity in the left and right lobes of the IC exhibited a small decrease at P10 (Figure 9E). To determine the spatial extent of neuronal activation in IC, we calculated the average area activated during an IC event (defined by pixels that exhibited an amplitude response greater than 2/3 of the maximum intensity; Figure 9F). The spatial spread of activity increased from P1 to P3, then slowly decreased over the next two postnatal weeks (Figure 9E). These results indicate that auditory neurons within isofrequency domains experience a prolonged period of correlated activity prior to hearing onset and that the domains of active neurons decrease with development, paralleling tonotopic refinement within the IC.

**Figure 9.**
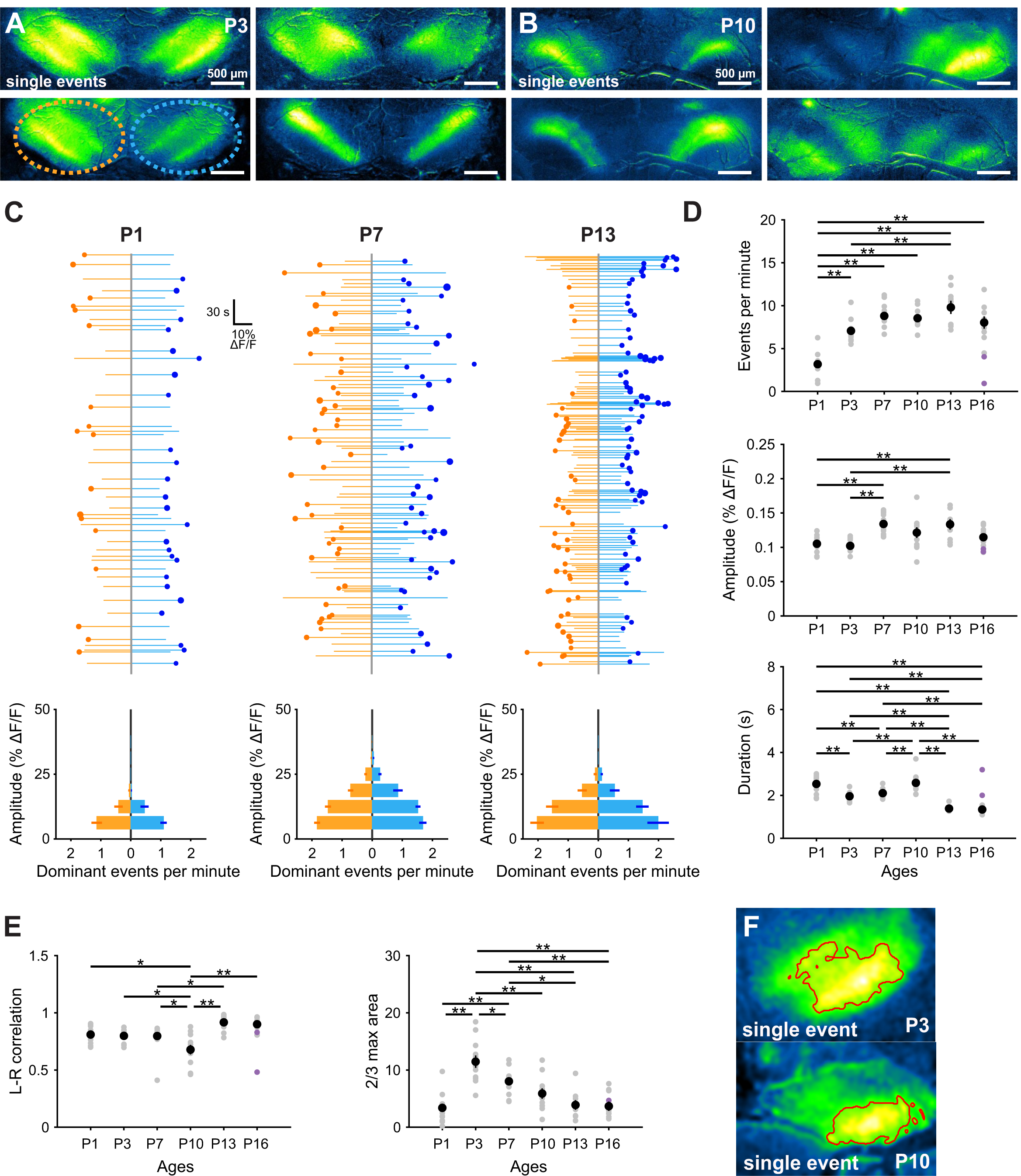
Developmental increase in spontaneous activity in the IC. (A) Images of spontaneous Ca^2+^ transients in the auditory midbrain (IC) of unanesthetized Snap25-T2A-GCaMP6s mice (P3). Orange and blue ovals indicate left and right IC, respectively, and correspond to activity traces in (C). (B) Images of spontaneous Ca^2+^ transients in the auditory midbrain (IC) of unanesthetized Snap25-T2A-GCaMP6s mice (P3). Note higher degree of lateralization compared than (A). (C) Graphs of activity over time for left (orange) and right (blue) lobes of the IC. Each line represents an individual event, the circle indicates which side had the greater intensity, and the size of dots represents the difference in fluorescence between the two sides. (bottom) Histograms showing the number of dominant events per amplitude bin. (D) Quantification of frequency, amplitude and duration (half-width of events) of events across different ages. n = 13 P1, n = 12 P3, n = 15 P7, n = 11 P10, n = 9 P13 and n = 9 P16 mice, (one-way ANOVA with Tukey post-hoc; **p < 0.005, comparisons not explicitly shown were not statistically significant). (E) Quantification of the left and right IC correlation coefficient (Pearson) and average area of each event (calculated as the area of pixels with values greater than 2/3 * max intensity) across different ages. (F) Example images of quantification of event areas (2/3 * max intensity delineated by red boundary) for single events.

### P2RY1 activity is required for spontaneous activity *in vivo* at P1

After the first postnatal week, spontaneous activity in the IC is sensitive to acute inhibition of P2RY1 (Babola et al., 2020). To determine whether P2RY1 is required for spontaneous activity in newborn animals, we performed time-lapse imaging of spontaneous activity in IC before and after acute injection MRS2500 (or vehicle) into the intraperitoneal space. Following MRS2500 injection, there was a significant decrease in the frequency (4.7 ± 0.9 events per minute in control, 2.49 ± 0.5 events per minute in MRS2500, Student’s t-test with Bonferroni correction, t(6) = 4.07, p = 0.01), but not the amplitude (0.077 ± 0.005 DF/F in control, 0.079 ± 0.005 DF/F in MRS2500, Student’s t-test with Bonferroni correction t(6) = 0.33, p = 0.76), of spontaneous Ca^2+^ transients (Figure 10D,F,G). In contrast, there were no significant changes in the frequency (Student’s t-test with Bonferroni correction t(7) = 0.52, p = 0.62) or amplitude (Student’s t-test with Bonferroni correction t(7) = 2.07, p = 0.08) after injection of vehicle (Figure 10B,E,G). Administration of MRS2500 did not alter the frequency or duration of retinal wave-induced activity in the SC (Figure 10G), suggesting the effects observed in IC are due to manipulation of P2RY1 receptors in the cochlea rather than CNS (Delekate et al., 2014). These *in vivo* results provide further evidence that P2Y1 autoreceptors within the cochlea initiate spontaneous bursts of neural activity in developing auditory centers from birth until the onset of hearing.

**Figure 10.**
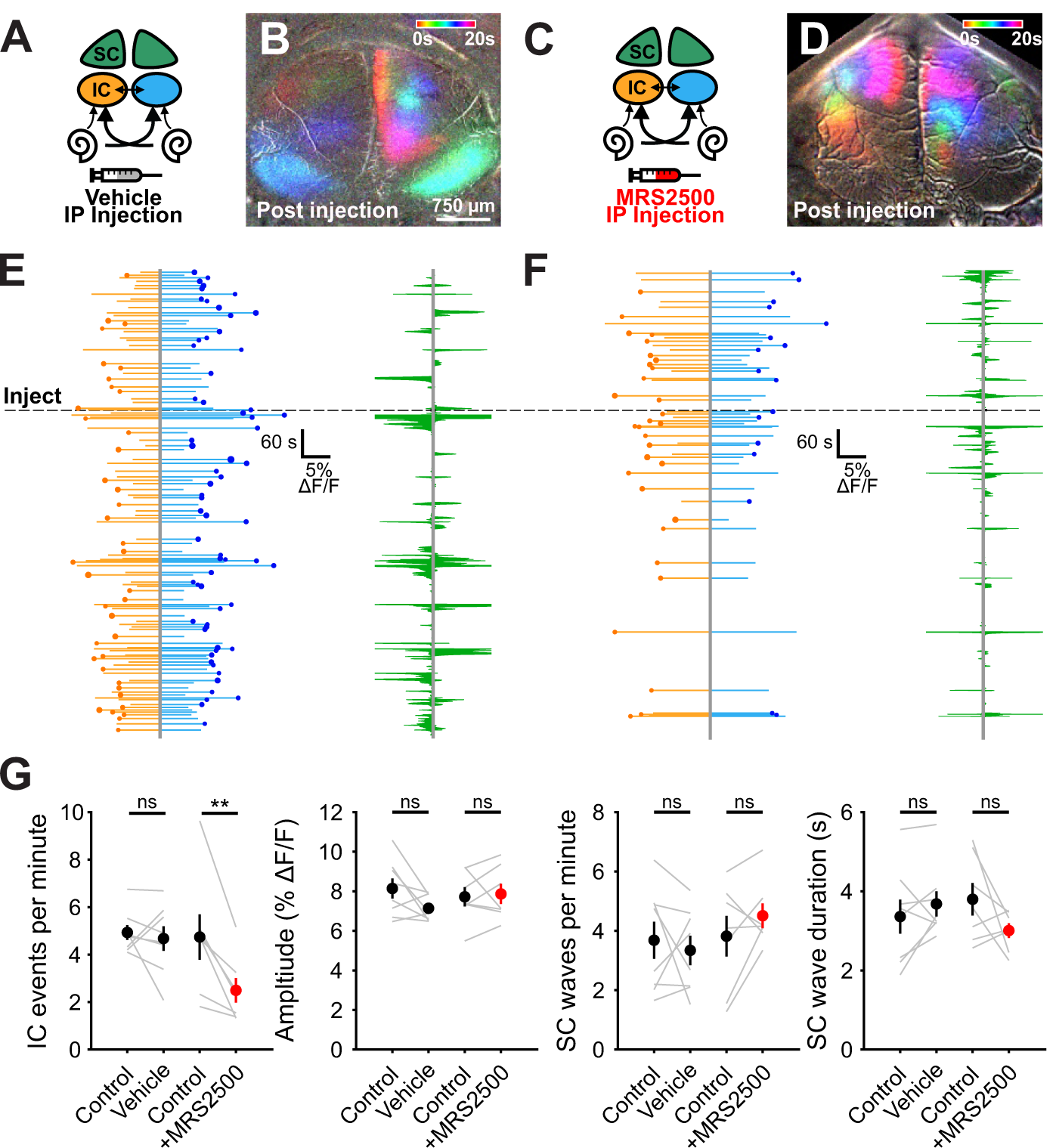
Early onset of spontaneous activity in the IC is P2RY1 dependent. (A) Schematic displaying flow of information from cochlea to midbrain. Sham solution (5% mannitol) was injected via IP catheter during imaging. (B) Ca^*2+*^ transients in the IC and SC after injection of sham solution. Transients are colored based on time of occurrence. segment after sham injection showing normal activity in both IC and SC. (C) Similar to (A), but with injection of MRS2500 via IP catheter during imaging. (D) Similar to (B), but following injection of MRS2500. (E-F) (left) Activity over time in left and right IC where each line indicated the fluorescence intensity of each detected event, the circle indicates the dominant lobe, and the size of the circle indicates the difference in fluorescence. Dashed line indicates time of injection. (right) SC activity before and after injection. Green shaded regions indicate the number of active ROIs in the left and right SC. (G) Quantification of IC event frequency and amplitude, and SC event frequency and duration. N = 8 sham injected and n = 7 MRS2500 injected *Snap25-T2A-GCaMP6s* mice, (paired t-test with Bonferroni correction applied, *p < 0.05, ns: not significant).

### Alpha 9-containing nicotinic acetylcholine receptors modulate bilateral activity patterns in IC

The results described above indicate that activation of P2RY1 on ISCs triggers a signaling cascade that coordinates the activity of nearby IHCs, SGNs, and central auditory neurons throughout the pre-hearing period. However, this is not the sole modulatory input to IHCs during this period. At this early stage of development, efferent cholinergic fibers form transient, inhibitory synapses on IHCs (Glowatzki and Fuchs, 2000), providing an additional means to shape IHC electrical activity. In *ex vivo* cochleae preparations, acute application of nicotinic acetylcholine receptor antagonists induces IHC burst firing, suggesting that release from cholinergic inhibition can initiate spontaneous bursts of activity (Johnson et al., 2011). Moreover, *in vivo* extracellular recordings from auditory brainstem neurons in anesthetized mice lacking the nicotinic acetycholine receptors in IHCs (Elgoyhen et al., 1994), exhibited bursts of action potentials at frequencies indistinguishable from controls, but bursts were shorter and contained more spikes (Clause et al., 2014), indicating that suppression of cholinergic inhibition of IHCs leads to altered burst firing of central auditory neurons. However, the influence of this efferent inhibitory input on the coordinated firing of auditory neurons *in vivo* in unanesthetized mice has not been examined. To explore the contribution of nAChRα9 signaling to macroscopic patterns of activity in IC, we performed time-lapse imaging of spontaneous activity from both nAChRα9 knockout (*α9* KO, *Chrnα9* ^*–/ –*^) and nAChRα9 gain-of-function (*α9* GOF; *Chrnα9*^L9’T/L9’T^ or *Chrnα9*^L9’T/+^) mice (Figure 11A-C and Supplemental Video 9), which exhibit prolonged efferent currents with slower desensitization kinetics in IHCs (Taranda et al., 2009; Wedemeyer et al., 2018). The frequency of spontaneous events in IC was unchanged in both *α9* KO and GOF mice relative to controls (One-way ANOVA, F(3,45) = 0.46, p = 0.71; Figure 11F). However, IC Ca^2+^ transients in homozygous *α9* GOF mice were unexpectedly larger in amplitude than controls; *α9* KO exhibited a trend towards lower amplitude Ca^2+^ transients, but this did not achieve significance (One-way ANOVA, F(3,45) = 18.22, p = 7E-8; Tukey HSD, p = 0.06; Figure 11F). These changes are opposite of what would be predicted from simply relieving or enhancing the inhibitory effect of acetylcholine on IHCs (Glowatzki and Fuchs, 2000). Similarly, individual events in homozygous *α9* GOF mice were longer (full width at half maximum) than controls (One-way ANOVA, F(3,45) = 3.2, p = 0.032; Tukey HSD, p = 0.026; Figure 11E,F), opposite of what would be predicted from greater inhibition of IHCs. There were also notable changes in the degree of lateralization among spontaneous IC events (Figure 11C, average event circles and Figure 11F, L-R correlation). Bilateral events were more symmetrical in *α9* GOF and less symmetrical in *α9* KO mice relative to controls (Figure 11F). Together, these results indicate that cholinergic efferent input to IHCs modulates the coordinated activity of central auditory neurons in unexpected ways, and influences interhemispheric representation of cochlear activity before hearing onset.

**Figure 11.**
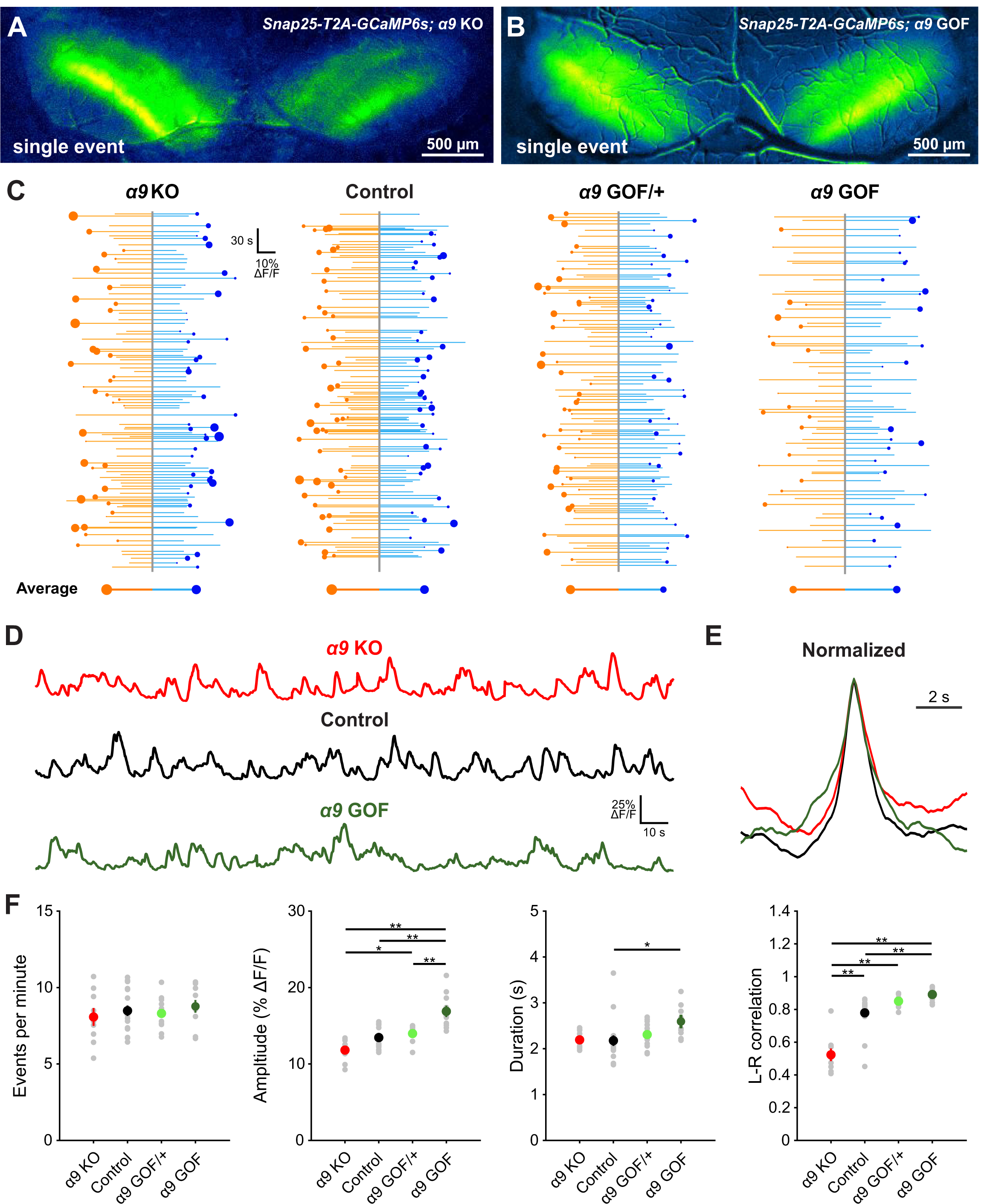
Cholinergic modulation of IHCs influences correlated activation of IC neurons before hearing onset. (A) Exemplar spontaneous Ca^2+^ transient in the auditory midbrain (IC) of unanesthetized *Snap25-T2A-GCaMP6s; Chrnα9* ^*–/–*^ (α9 KO) mice (P7). (B) Exemplar spontaneous Ca^2+^ transient in the auditory midbrain (IC) of unanesthetized *Snap25-T2A-GCaMP6s; Chrnα9*^*L9’T/L9’T*^ (α9 GOF) mice (P7). (C) Graphs of activity over time for left (orange) and right (blue) lobes of the IC for indicated genotypes. Each line represents an individual event, the circle indicates which side had the greater intensity, and the size of dots represents the difference in fluorescence between the two sides. (bottom) Average event for each lobe of the IC. Size of circle is the average difference in the fluorescence between the two sides. (D) Example fluorescence traces for indicated genotypes. (E) Average event from traces shown in (D) normalized to amplitude. (F) Quantification of event frequency, amplitude, duration and left-right correlation coefficient (Pearson) across indicated genotypes. n = 9 α9 KO, n = 17 control, n = 13 α9 GOF/+, and n = 10 α9 GOF mice, (one-way ANOVA with Tukey post-hoc; **p < 0.005, *p < 0.05, comparisons not shown are not statistically significant).

## DISCUSSION

### Generation of coordinated neural activity by cochlear supporting cells

Nascent neural networks in the developing CNS exhibit highly stereotyped spontaneous activity, consisting of periods of high frequency action potential firing interspersed with long periods of quiescence (Blankenship and Feller, 2010). Similar to the visual system, spontaneous activity generated in the cochlea begins just prior to birth in mice (Figure 1), providing a prolonged period over which activity-dependent maturation and refinement can occur before the middle ear opens (∼P12); however, much less is known about the mechanisms that drive spontaneous activity or how this activity changes over this developmental period. In contrast to the dynamic mechanisms responsible for retinal wave generation, our studies indicate that bursts in the auditory system are driven consistently by ISC purinergic signaling. Based on measures of spontaneous activity *in vivo*, each auditory neuron will experience more than 30,000 discrete bursts (∼2.0 bursts/minute; ∼2900 bursts/day) prior to hearing onset (Tritsch et al., 2010; Clause et al., 2014; Babola et al., 2018). Consistent with the stable generation of P2RY1-dependent bursts, neural activity in the IC remained highly stereotyped during the first two postnatal weeks, providing a means for activity-dependent, Hebbian plasticity during this early developmental period.

### Purinergic signaling in the developing and adult cochlea

Despite widespread expression of ionotropic P2X and metabotropic P2Y receptors in the developing cochlea (Nikolic et al., 2003; Lahne and Gale, 2008; Huang et al., 2010; Liu et al., 2015; Wang et al., 2020), ISC electrical activity and highly structured burst firing of SGNs appears reliant primarily on P2RY1. The lack of P2X or other G_q_-coupled P2Y receptor activation may reflect the temporal and spatial characteristics of ATP release, which may occur in locations abundant in P2RY1 or yield ATP metabolites that favor P2RY1 activation. However, burst firing persists in *P2ry1* KO mice, in which IHCs are more depolarized (Babola et al., 2020), perhaps reflecting compensatory or activation of other purinergic receptors that are normally subthreshold.

A similar diversity of purinergic receptor expression is observed in the adult cochlea, including P2X receptors, metabotropic P2Y, and adenosine P1 receptors (Housley et al., 2009; Huang et al., 2010). ATP receptor activation appears to play a neuroprotective role, as endolymphatic ATP increase following trauma and infusion of ATP into the inner ear profoundly reduces sound-evoked compound action potentials in the auditory nerve. While these effects may reflect shunting inhibition through P2×2 receptors (Housley et al., 2013), recent evidence indicates that supporting cells in the mature cochlea continue to exhibit large Ca^2+^ transients in response to exogenous ATP and UTP (Zhu and Zhao, 2010; Sirko et al., 2019). The role of this activity is unclear, but Ca^2+^ transients induced by mechanical damage and subsequent ATP release trigger ERK1/2 activation and promote IHC death in the developing cochlea (Lahne and Gale, 2008), limiting excitotoxic damage. If the developmental pathways described here reemerge following traumatic noise damage, purinergic receptor signaling could enhance K^+^ redistribution, reduce IHC depolarization and reduce excitotoxic damage.

### The emergence of ATP induced extracellular space changes in the cochlea

ISCs crenations dramatically increase the volume of extracellular space and speed K^+^ redistribution, shaping the envelope of IHC excitation following P2RY1 activation (Babola et al., 2020). Prior studies demonstrated that crenations were associated with only the largest increases in Ca^2+^ (Tritsch et al., 2007), perhaps reflecting the activation range of TMEM16A channels. However, ISCs in P0 cochleae exhibited spontaneous currents as large as those observed at P7, and yet ISCs at P0 did not exhibit crenations. It is possible that ion flux in ISCs at P0 is lower, reducing the osmotic force. *P2ry1* promoter activity is lower at P0 (Figure 2), TMEM16A protein levels increase during the first postnatal week (Wang et al., 2015), and both *P2ry1* and TMEM16A mRNA expression rapidly increase over the first postnatal week (Scheffer et al., 2015; Kolla et al., 2020). Accompanying these expression changes, the charge transfer of all events was similar across the first two postnatal weeks, despite decreasing event frequency (Figure 1D), indicating a moderate increase in ion flux/event over development, enhancing osmotic force during each event.

While changes in ion flux may contribute to the emergence of crenations during the first postnatal week, expression of aquaporins, a family of highly permeable water channels that enable rapid diffusion of water across biological membranes (Reuss, 2012), may also regulate this process. Recent single-cell RNAseq analysis of the cochlear epithelium revealed that AQP4 and AQP11 genes are expressed at similar levels within Kölliker’s organ after the first postnatal week (Kolla et al., 2020). AQP11 is a non-traditional aquaporin family member, sharing only 20% homology with other aquaporin family members (Ishibashi et al., 2000), is localized to the ER membrane (Morishita et al., 2005), has a higher permeability to glycerol than water (Madeira et al., 2014), and is expressed at relatively stable levels throughout cochlear development (Kolla et al., 2020), making it an unlikely candidate to regulate water movement into the extracellular space. AQP4 is highly permeable to water and its expression dramatically increases between P1 and P7 (Kolla et al., 2020), indicating that a developmental increase in AQP4 expression could enable the large movements of water that underlie ISC crenation.

Recent evidence suggests that ISC control of the extracellular space influences the activation of IHCs during spontaneous events and controls IHC excitability. Conditional removal of TMEM16A from the cochlea (Wang et al., 2015) or acute inhibition of P2RY1 (Babola et al., 2020) both prevent ISC crenation during spontaneous events. This collapse of the extracellular space limits K^+^ diffusion, reducing the number of IHCs activated per ISC Ca^2+^ transient and promoting local K^+^ buildup that eventually leads to tonic IHC firing. Therefore, the lack of crenation at birth may slow K^+^ redistribution to induce larger and more prolonged depolarization of IHCs at a time when ribbon synapses are immature and Ca^2+^ channel expression is low (Marcotti, 2012; Michanski et al., 2019).

### Involvement of cholinergic efferents in modulating early spontaneous activity

In the developing cochlea, IHCs are transiently innervated by efferent fibers, which provide powerful inhibitory input (Glowatzki and Fuchs, 2000). Previous studies in isolated cochleae demonstrated that temporarily relieving this inhibition produced bursts of action potentials in IHCs (Johnson et al., 2011), suggesting that efferent activity may initiate burst firing. However, auditory brainstem neurons in (anesthetized) α9 KO mice exhibit prominent burst firing *in vivo*, with bursts occurring at similar frequencies, but with shorter durations and containing more action potentials than controls (Clause et al., 2014). Moreover, α9 GOF mice that have enhanced efferent inhibition of IHCs (Wedemeyer et al., 2018), also exhibit spontaneous action potentials in the auditory brainstem, although at lower frequencies than controls (Di Guilmi et al., 2019). Our *in vivo* macroscopic imaging studies indicate that periodic excitation of auditory midbrain neurons occurred in α9 KO mice at the same frequency as controls (Figure 11), reinforcing the conclusion that efferent input is not required to initiate burst firing. However, manipulating nAChRα9 signaling altered the lateralization (contralateral bias) of activity across the brain. In α9 KO mice, bilateral activation of the IC was more asymmetric than controls (Figure 11A,F), and conversely, activity in α9 GOF mice was more symmetric across the IC (Figure 11B,F). Bilaterally symmetric events reflect activity initiated in one cochlea (Babola et al., 2018); thus, these changes in propagation of activity to both hemispheres may arise through alterations in the precise patterning of firing (Clause et al., 2014; Di Guilmi et al., 2019) due to changes in the electrophysiological properties of auditory neurons (Di Guilmi et al., 2019). Surprisingly, the effects on IC neuronal Ca^2+^ transients in α9 KO and GOF mice were opposite of that predicted based on the inhibitory effect of acetylcholine on IHCs (Glowatzki and Fuchs, 2000), with enhanced inhibition of IHCs resulting in larger amplitude events in central auditory neurons (Figure 11). The differential effects of manipulating efferent inhibition on spontaneous activity as it propagates through the auditory system (Di Guilmi et al., 2019), raise the possibility that specific patterns of activity are required to induce proper maturation of sound processing circuits.

### The role of spontaneous neural activity in development

Silencing thalamic input to the somatosensory cortex during early postnatal life results in failure of barrel field formation (Antón-Bolaños et al., 2019) and genetic disruption of retinal waves leads to profuse retinal ganglion cell axon arborization in SC and segregation-deficits in the thalamus (Rossi et al., 2001), demonstrating the critical role of early patterned activity in circuit maturation. In the auditory system, similar refinement deficits have been observed in α9 KO mice and in various models of deafness. However, the functional consequences of these manipulations on sensory performance remain underexplored. α9 KO mice exhibit deficits in sound localization tasks (Clause et al., 2017), but whether these changes are due to disruption in spontaneous activity or the lack of a functional efferent system after hearing onset remains uncertain. Insight into the mechanisms that govern spontaneous activity in the auditory system provide an experimental framework for selectively disrupting early spontaneous activity, while preserving hearing, allowing assessment of the role of stereotyped burst firing in development.

## ACKNOWLEDGEMENTS

We thank Dr. M Pucak and N Ye for technical assistance, T Shelly for machining expertise, and members of the Bergles laboratory for discussions and comments on the manuscript. TB was supported by an NRSA from the NIH (F31DC016497). Funding was provided by grants from the NIH (DC008060, NS050274 to DB, DC001508 to ABE), the Mathers Foundation (Grant #MF-1804-00065), and the Rubenstein Fund for Hearing Research.

## SUPPLEMENTAL VIDEO LEGENDS

**Supplemental video 1. Delayed onset of P2ry1-dependent spontaneous crenations in ISCs**.

(A) DIC imaging of spontaneous cell shrinkage events (crenation) in the cochlea across the prehearing period. Application of MRS2500 (1 μM), a selective P2RY1 antagonist, is indicated in the top right corner of each video.

**Supplemental video 2. Grid-based analysis of Ca**^**2+**^ **transients**.

(A) Time lapse imaging of isolated cochlea from P0 *Pax2-Cre;R26-lsl-GCaMP3* mice. White squares indicate active ISCs and white dots indicate active IHCs.

**Supplemental video 3. Correlated activation of IHCs and ISCs requires P2RY1 signaling**.

(A) Time lapse imaging of isolated middle sections of cochlea from P0 *Pax2-Cre;R26-lsl-GCaMP3* mice. Application of MRS2500 (1 μM) is indicated in the top right corner of each video.

**Supplemental video 4. Tonotopic differences in extent of IHC activation at early developmental time points**.

(A) Time lapse imaging of isolated apical and basal sections of cochlea from *Pax2-Cre;R26-lsl-GCaMP3* mice (P0).

**Supplemental video 5. Tonotopic differences in extent of SGN activation at early developmental time points**.

(A) Time lapse imaging of SGN activity in isolated apical and basal sections of cochlea from E16.5 and P0 *Snap25-T2A-GCaMP6s* mice.

**Supplemental video 6. Correlated activation of SGNs requires IHC glutamate release**.

(A) Time lapse imaging of SGN activity in isolated basal sections of cochlea from P0 *Snap25-T2A-GCaMP6s* mice. Application of CNQX/CPP (50/100 μM) is indicated in the top right corner.

**Supplemental video 7. Correlated activation of SGNs requires P2ry1-mediated excitation of IHCs**.

(A) Time lapse imaging of SGN activity in isolated basal sections of cochlea from P0 *Snap25-T2A-GCaMP6s*. Application of MRS2500 (1 μM) is indicated in the top right corner.

**Supplemental video 8. Developmental increase in spontaneous activity in the IC**.

**(A)** Time lapse imaging of inferior colliculus activity in unanesthetized *Snap25-T2A-GCaMP6s* mice.

**Supplemental video 9. Cholinergic modulation of IHCs influences correlated activation of IC neurons before hearing onset**.

(A) Time lapse imaging of IC activity in unanesthetized, P7 *Snap25-T2A-GCaMP6s; Chrnα9* ^*–/–*^ (*α9* KO) and *Chrnα9* ^*L9’T/L9’T*^ (*α9* GOF) mice.

